# Computational mechanisms for neural representation of words

**DOI:** 10.1101/2021.06.27.450093

**Authors:** Ze Fu, Xiaosha Wang, Xiaoying Wang, Huichao Yang, Jiahuan Wang, Tao Wei, Xuhong Liao, Zhiyuan Liu, Huimin Chen, Yanchao Bi

**Author notes:** Correspondence to Y. Bi.

## Abstract

A critical way for humans to acquire, represent and communicate information is through language, yet the underlying computation mechanisms through which language contributes to our word meaning representations are poorly understood. We compared three major types of word computation mechanisms from large language corpus (simple co-occurrence, graph-space relations and neural-network-vector-embedding relations) in terms of the association of words’ brain activity patterns, measured by two functional magnetic resonance imaging (fMRI) experiments. Word relations derived from a graph-space representation, and not neural-network-vector-embedding, had unique explanatory power for the neural activity patterns in brain regions that have been shown to be particularly sensitive to language processes, including the anterior temporal lobe (capturing graph-common-neighbors), inferior frontal gyrus, and posterior middle/inferior temporal gyrus (capturing graph-shortest-path). These results were robust across different window sizes and graph sizes and were relatively specific to language inputs. These findings highlight the role of cumulative language inputs in organizing word meaning neural representations and provide a mathematical model to explain how different brain regions capture different types of language-derived information.

## Introduction

A typical adult human brain stores the forms and meanings of approximately tens of thousands of words (42,000 for typical English-speaking Americans) (Brysbaert, Stevens, Mandera, & Keuleers, 2016), which allows naming objects and actions, understanding and producing sentences, and contributing to various kinds of reasoning. Decades of neuroimaging and neuropsychological literature have studied the cognitive neural representations of word meaning and revealed that they are (at least partly) derived from sensory experiences distributed across multiple sensory association cortices (Fernandino et al., 2016; A. Martin, 2016; Miceli et al., 2001), with those sharing physical properties (sensory/motor experiences) represented more closely in corresponding brain regions (Binder et al., 2016; Clarke & Tyler, 2014, 2015). The roles of the language system in representing word meaning have recently been highlighted, with nonsensory, language-derived representation of word meaning (i.e., color and other visual concepts in congenital blindness) being identified in the dorsal part of the anterior temporal lobe (ATL) (Striem-Amit, Wang, Bi, & Caramazza, 2018; Xiaoying Wang, Men, Gao, Caramazza, & Bi, 2020). This ATL cluster, along with the left lateralized frontal-temporal cortices, including the inferior frontal gyrus (IFG) and the posterior part of the middle temporal gyrus, also show stronger sensitivity to abstract words than concrete words (Binder, Desai, Graves, & Conant, 2009; Hoffman, Binney, & Ralph, 2015; Noppeney & Price, 2004; J. Wang, Conder, Blitzer, & Shinkareva, 2010), indicating its potential role in representing meaning representation derived from language.

Does the brain capture word meaning representation from cumulative language experience, and if yes, how? It has recently been shown that word relations constructed from various natural language computation models derived from a large text corpus correlate with the fMRI whole-brain response patterns to corresponding words (count models (Huth, De Heer, Griffiths, Theunissen, & Gallant, 2016; Mitchell et al., 2008); GloVe model (Anderson et al., 2019; Pereira et al., 2018)) and with the brain activity patterns of more specialized language processing regions/networks (word2vec and LSA models (Carota, Kriegeskorte, Nili, & Pulvermüller, 2017; Carota, Nili, Pulvermüller, & Kriegeskorte, 2021; Xiaosha Wang et al., 2018)). These observed correlations between language computation models and brain activity patterns suggest that word representation in the brain may follow certain types of computation from language inputs, but their interpretation requires caution. There are significant correlations between word-relational structures constructed from different language computation models, as well as with relational structures derived from the sensory systems (e.g., “cat” and “dog” are closely related across all of these different types of measures) (Kumar, 2021; Lenci, 2018). Understanding the underlying computational architecture (and potentially algorithms) of the language-derived knowledge system requires carefully examining the differences in these models and testing which models better explain the brain activity patterns associated with word meanings.

Ample evidence has shown that humans are sensitive to various types/scales of statistical patterns from language-like inputs (Lynn & Bassett, 2020). Statistical learning studies show that humans can detect both variations in the probabilities of transitions between items and in non-local network structures (e.g. non-adjacent dependency, community structures) (Garvert, Dolan, & Behrens, 2017; Saffran, Aslin, & Newport, 1996; Schapiro, Rogers, Cordova, Turk-Browne, & Botvinick, 2013). Accordingly, for the statistical patterns derived from the long-term representation of the cumulative language inputs, we consider three major types of computation mechanisms: 1) Simple co-occurrence, which involves the collective counts or probabilities of two words co-occurring in a context window. Simple co-occurrence measures, reflecting the first-order proximity between two words, have been used to model word relatedness (Church & Hanks, 1990; Recchia & Jones, 2009; Spence & Owens, 1990). 2) Graph representation based on raw simple co-occurrence patterns (Borge-Holthoefer & Arenas, 2010; Jackson & Bolger, 2014), with different types of relationships/distances computed from the graph-related structure (e.g. common neighbors, shortest path). It has been shown that different types of word relations (similarity and association) can be measured using different graph measures implemented in a unified word co-occurrence graph space, providing a relatively transparent, dissociable representation of word statistical relation patterns (Jackson & Bolger, 2014). 3) Vector representation based on a set of fixed dimensions obtained through matrix factorization methods (e.g. LSA) (Landauer & Dumais, 1997) or model-based embedding methods (e.g. word2vec, GloVe) (Mikolov, Chen, Corrado, & Dean, 2013; Pennington, Socher, & Manning, 2014). These vector models, especially embedding models (word2vec, GloVe), have also been shown to achieve state-of-the-art performance in a variety of semantic-related evaluation tasks (Baroni, Dinu, & Kruszewski, 2014; Pereira, Gershman, Ritter, & Botvinick, 2016). While these vector models are computationally efficient and widely used in modeling behavioral and neural semantic representations, the statistical information embedded in the obtained vector space (e.g., the neural network embedding one) has remained elusive due to explicit/implicit dimension reduction and hyperparameter tuning processes (Levy & Goldberg, 2014; Levy, Goldberg, & Dagan, 2015).

Here, we compare these three major types of computation mechanisms that capture different aspects of statistical patterns of the language corpus in modeling neural activity patterns of word processing: simple co-occurrence, graph-space relations and neural-network based vector-space relations (i.e., cosine distance in a learnt word2vec space) (Fig. 1). We conducted a word production fMRI experiment (oral picture naming) and a word recognition fMRI experiment (word familiarity judgment) on the same set of 95 words to obtain their neural activity patterns across different input/output modalities. Distances calculated from simple co-occurrence, graph-related measures (graph-common-neighbors and graph-shortest-path) from a large-scale language corpus (Chinese Web n-gram Corpus; https://catalog.ldc.upenn.edu/LDC2010T06) (Liu, Yang, & Lin, 2010) and vector cosine distance from a pretrained, open-accessed word2vec models (Li et al., 2018) were calculated for these 95 words. Using representation similarity analysis (RSA) (Kriegeskorte, Mur, & Bandettini, 2008), the representational dissimilarity matrices (RDMs) computed from these different language computation models were fit with the neural RDMs derived from the fMRI data to locate neural circuits that are organized by specific language statistical properties. Graph-space relations of visual co-occurrence statistics derived from a large visual image database (VisualGenome; https://visualgenome.org/) (Krishna et al., 2017) were also compared to examine the universality/specificity of the observed word neural computations.

**Fig. 1.**
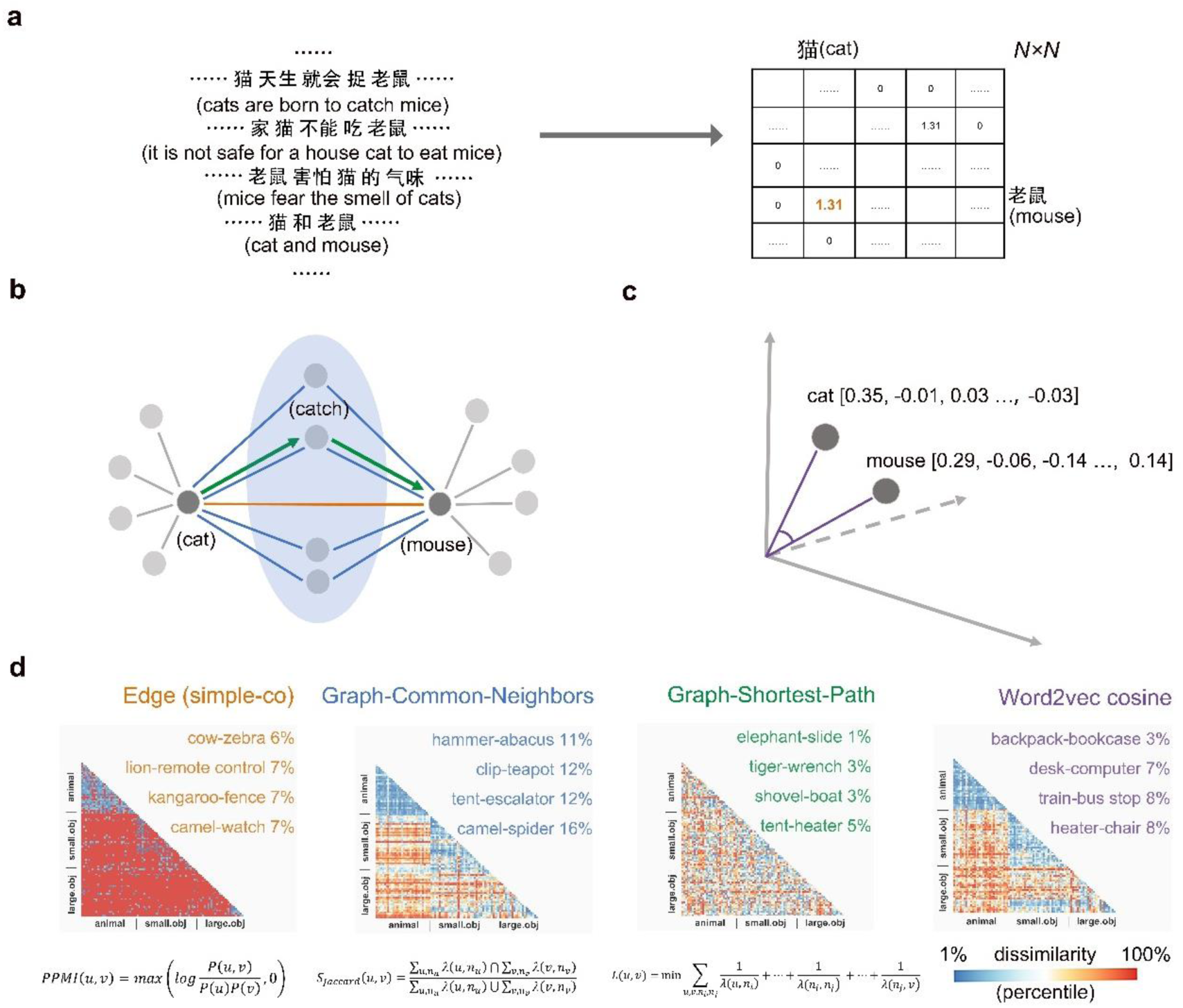
Construction of language computation models for word (meaning) representation. (**a**) Word simple co-occurrence statistics derived from cumulative language inputs. The raw co-occurrence counts in a given text “context” (window size is 2 in the bigram version) were obtained to construct a word co-occurrence matrix (from a downsampled Chinese Web Google n-gram Corpus, including 83,007 unique word samples). The raw co-occurrence counts were normalized into PPMI values to represent simple co-occurrence in the long-term language experience. (**b**) Graph representation and related measures. The N by N word co-occurrence matrix was transformed into a graph representation, where each unique word was treated as a node and PPMI values between word pairs as edges (N = 83,007, with 34,586,840 word co-occurrence relationships). Two kinds of graph-related measures were calculated in the graph space: common neighbors and shortest-path distances. (**c**) Neural-network-based vector embedding representation. A pretrained and open-accessed word2vec model was selected (Li et al., 2018). In the word2vec model, words were represented as a 300- dimensional vector, and word relations were computed as the cosine distance between these vectors. (**d**) Results of language RDMs. Four types of language-derived computation RDMs from three major representations mentioned above were measured (edge RDM (simple co-occurrence), graph-CN RDM, graph-SP RDM and word2vec RDM). The value of dissimilarity was transformed to a percentile for display. Red indicates high dissimilarity (low similarity), and blue indicates low dissimilarity (high similarity). Four sample word pairs with strong association relationships (low distance score) were presented, which rank in the top n (the exact number of n presented at the right side of word pairs) in one measure and the bottom 50% in the other three measures. Mathematical formulas for edge model and each graph-related model were presented below (see Methods for detailed information).

## Results

### Construction of language-computation-model-based RDMs

Three types of language computation mechanisms were implemented. Simple co-occurrence counts were derived from a large-scale language corpus (Chinese Web n-gram Corpus, consisting of approximately 883 billion Chinese words). These raw counts were PPMI-normalized to represent first-order proximity between two words (Fig. 1a). Such normalized word co-occurrence of the 95 experimental stimuli (Fig. S1 & Table S1) was used to construct the 95 x 95 simple co-occurrence RDM. Beyond the simple co-occurrence (i.e. edge RDM in the graph space), graph-common-neighbors (graph-CN) and graph-shortest-path (graph-SP), two types of graph-related measures reflecting different aspects of statistical properties were computed in a downsampled graph space (83,007 unique Chinese word samples as nodes, 34,586,840 PPMI-normalized simple co-occurrence as edges) to yield a graph-CN RDM and a graph-SP RDM (Fig. 1b). A word2vec RDM was constructed based on the cosine distance in a state-of-art pretrained word vector dataset (Li et al., 2018) (Fig. 1c). Visualizations of these four computational RDMs are presented in Fig. 1d. Spearman rank correlation analysis revealed that these RDMs were moderately to highly intercorrelated (edge RDM with CN RDM, Spearman *r* = 0.48; edge RDM with SP RDM, Spearman *r* = 0.40; edge RDM with word2vec RDM, Spearman *r* = 0.45; CN RDM with SP RDM, Spearman *r* = 0.53; CN RDM with word2vec RDM, Spearman *r* = 0.68; SP RDM with word2vec RDM, Spearman *r* = 0.40; *Ps* < 1.29 x 10^-172^).

### RSA results: Relationship between language models and brain activity patterns

Neural RDMs of the 95 items were generated and fit with language model RDMs for each fMRI experiment in an iterative sphere (10 mm) of each individual native space (an individually defined gray matter mask), following the procedure of whole brain searchlight RSA (Kriegeskorte, Goebel, & Bandettini, 2006) (Fig. 2a). In each experiment, analyses were carried out for stimuli peripheral variables to perform sanity checks: pixel RDM and gist RDM in the oral picture naming experiment, and pixel RDM and familiarity (button-press) RDM in the word familiarity judgment experiment. The RSA results of these control models were highly consistent with the previous literature (Carota et al., 2021; Devereux, Clarke, Marouchos, & Tyler, 2013; Kriegeskorte, Mur, Ruff, et al., 2008) (Fig. S2). In the main analyses of the language computation models, we first looked at the RSA results of each model independently (“raw” effect), with the peripheral factors (pixel and gist RDM in oral picture naming, pixel and word familiarity RDM in word familiarity judgment) regressed out. Given that these RDMs of language computation models are correlated (Fig. 1d), we further carried out a “unique effect” RSA for each language computation model, where the effects of the other language models were further controlled for (all using partial Spearman rank correlations). The convention cluster extent-based inference threshold (primary voxelwise *p* < 0.001, FWE-corrected cluster-level *p* < 0.05) was adopted. The results for both fMRI experiments are shown, with positive results across both experiments, i.e., the shared cognitive components (word meanings) across experimental inputs/outputs, presented in detail (Table 1). As the conjunction methods are relatively conservative (Nichols, Brett, Andersson, Wager, & Poline, 2005), we reported clusters that survived from the threshold (uncorrected voxelwise *p* < 0.005, cluster size > 20 voxels) across two experiments.

**Fig. 2.**
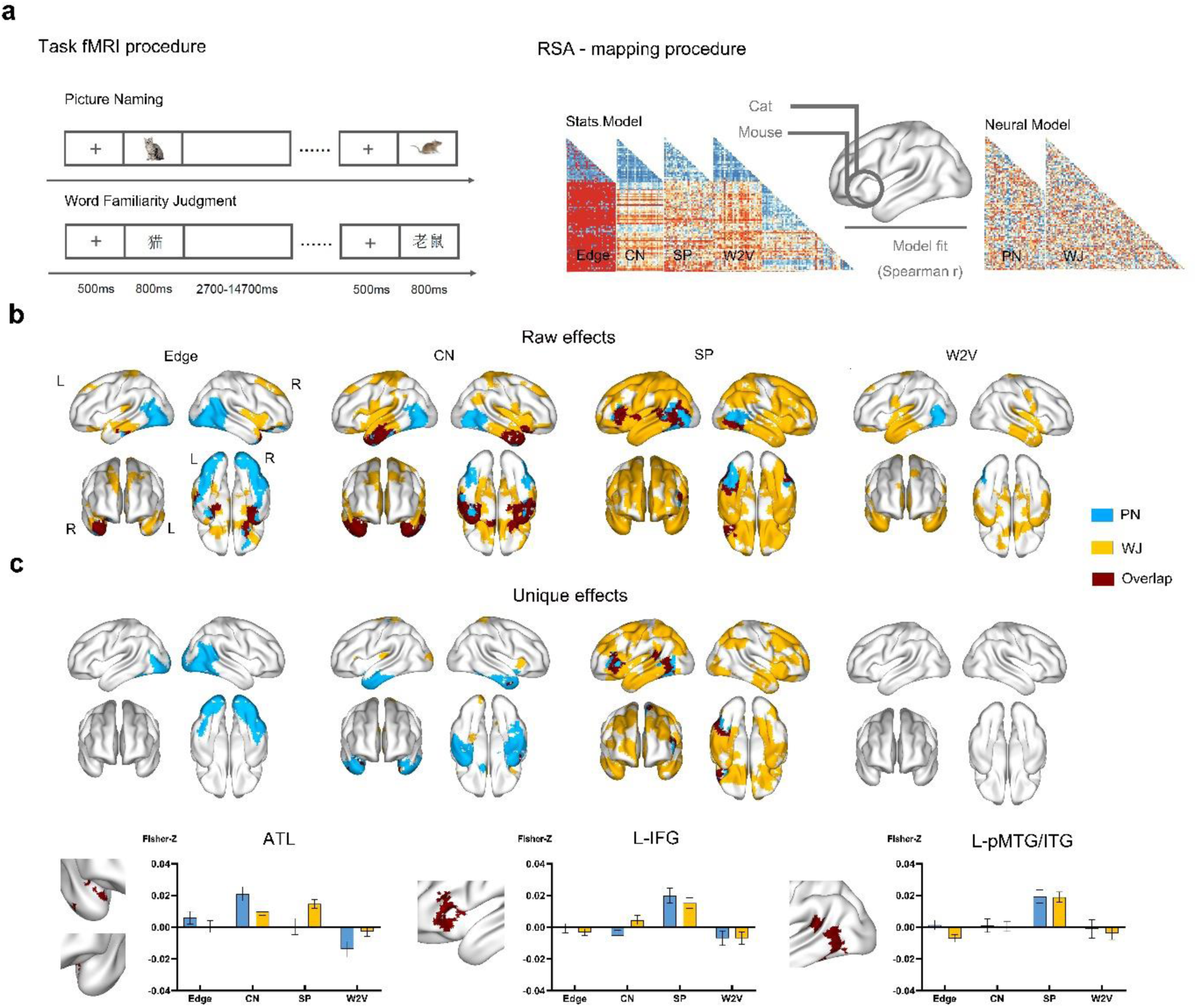
Overlap results of whole brain searchlight RSA across two types of word processing experiments revealed task-invariant neural effects on language graph-related measures in verbal sensitive regions. (**a**) Task fMRI design and RSA mapping procedure. Neural responses of 95 items under two types of word processing experiments (oral picture naming and word familiarity judgment) were collected using condition-rich event-related design. Pair-wise dissimilarities of these 95 neural activity patterns were used to construct neural RDMs. Second-order correlations between language computation RDMs (simple co-occurrence (edge) RDM, graph-CN RDM, graph-SP RDM and word2vec RDM) and neural activity patterns within a searching sphere (10 mm) were calculated. A significant Spearman *r* value served as an indicator that the information captured by the model is represented in the brain. (**b**) Overlap results of the raw effects of different language computation models across two types of word processing experiments (uncorrected voxelwise p < 0.005, cluster size > 20 in each type of experiment. (**c**) Overlap results of the unique effects (regressing the other three) across two types of word processing experiments. Graph-related distance showed a unique explanatory power in capturing neural activity patterns of verbal sensitive regions (graph-common-neighbors distance in bilateral ATL and graph-shortestpath distance in left IFG and pMTG/ITG). The bar plots for Fisher-Z transformed *r* values in the overlapping regions are presented for visualization without additional statistical inference. Clusters located in bilateral ATL (right aMTG, TP and left aPHG) were treated as one region, as results in each cluster yields similar patterns.

**Table 1.**
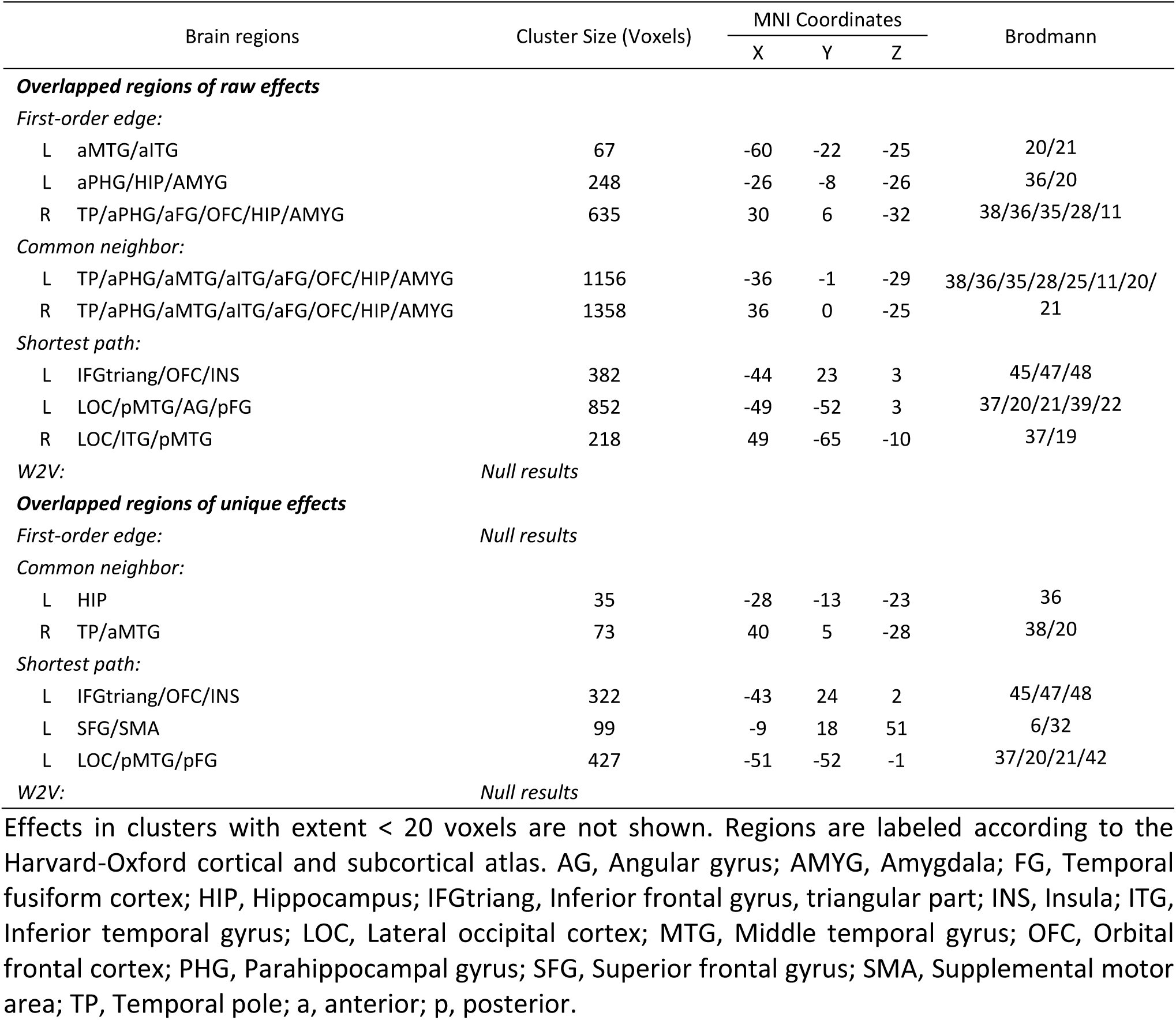
Overlap results of language computation model RSA across two types of word processing experiments.

#### Language model-brain RSA raw results

The maps of group-level whole brain searchlight RSA results from each language computation model in each experiment are shown (Fig. 2b; more detailed presentation for oral picture naming experiment in Fig. S3a and Table S2, for word familiarity judgment experiment in Fig. S3b and Table S3).

##### First-order-edge (simple co-occurrence) distance

In the oral picture naming experiment, the edge RDM correlated significantly with neural RDMs throughout the bilateral occipital-temporal cortex, with peak effects in the lateral occipital cortex (LOC), extending into the early visual cortex, pMTG, and the posterior division of the temporal fusiform gyrus (pFG) and bilateral ATL, including the right temporal pole (TP), anterior division of the temporal fusiform gyrus (aFG) and parahippocampal gyrus (aPHG). In the word familiarity judgment experiment, the neural effects of edge RDM were confined to the bilateral ATL, including the TP, aPHG and left anterior division of the middle temporal gyrus (aMTG), the dorsal part of the medial frontal cortex (medPFC), orbital frontal cortex (OFC), right precuneus, precentral and postcentral gyrus, cingulate gyrus, insula and putamen. The overlap analysis showed that the bilateral ATL, especially the ventral part, was sensitive to the edge RDM in a task-invariant manner.

##### Graph-common-neighbors distance

In the oral picture naming experiment, neural effects of graph-CN RDM were also found in occipital and temporal regions, including the bilateral LOC, pMTG, pFG and ATL. Clusters in the bilateral ATL encompassed the TP, aPHG, aFG, aMTG and inferior temporal gyrus (aITG). In the word familiarity judgment experiment, similar patterns were found in regions of bilateral ATL. Clusters extended into the medial temporal fusiform gyrus (medFG), orbital frontal cortex (OFC) and subcortical regions, including the hippocampus, amygdala, caudate and thalamus. More dorsally, clusters were found in the medPFC, precuneous cortex, cingulate gyrus, precentral gyrus and postcentral gyrus, right angular gyrus (AG), posterior part of the STG, bilateral insular cortex and primary auditory cortex. The overlap analysis showed the task invariant neural representation of the graph-CN RDM in the bilateral ATL, including the TP, aFG, aPHG, aMTG and aITG, the OFC and the subcortical regions, including the hippocampus and amygdala.

##### Graph-shortest-path distance

In the oral picture naming experiment, the neural activity patterns in the frontal-temporal cortex were significantly associated with the graph-SP RDM, including the bilateral LOC, pFG, pMTG, left insula and the pars triangularis part of the left IFG. More robust results were found in the word familiarity judgment experiment, which spread across the distributed brain regions, including bilateral temporal regions (with peak effects located in the left aSTG, aMTG, left amygdala), frontal regions (with peak effects located in the bilateral OFC, right caudate and right medPFC) and widespread clusters located in the parietal cortex. The overlap analysis revealed that the task-invariant representation of the graph-SP RDM was located in the bilateral LOC, bilateral pMTG, left pFG, ventral part of the left AG and pars triangularis part of the left IFG, as well as the OFC and insula.

##### Word2vec cosine distance

In the oral picture naming experiment, the neural patterns of bilateral LOC extending into pMTG were found to be significantly correlated with word2vec RDM. In the word familiarity judgment experiment, significant mapping between the word2vec RDM and neural RDMs was found in the bilateral ATL, extending into the medFG, OFC and subcortical regions, including the hippocampus, putamen and right thalamus. More dorsally, clusters were found in the precentral and postcentral gyrus, as well as the insula and primary auditory cortex. No voxels survived in the overlap analysis when investigating the task-invariant neural representation of word2vec RDM.

#### Language model-brain RSA unique results

The unique RSA effects of each language computation model, with the effects of other language models (and peripheral variables) partially removed, reveal the relative specificity of the target model in explaining the neural activity in a particular brain region (see above). The results surviving the conventional cluster extent-based inference threshold were presented in each experiment separately (oral picture naming in Fig. S3a and Table S2; word familiarity judgment in Fig. S3b and Table S3). The overlap results across the two experiments are shown in Fig. 2c and Table 1.

##### First-order-edge (simple co-occurrence) distance

In the oral picture naming experiment, the unique neural effects of edge RDM were located in the bilateral LOC, pFG and early visual cortex. In the word-judgment experiment, no regions showed significant effects.

##### Graph-common-neighbors distance

The significant mappings between the graph-CN RDM and neural RDMs in the bilateral ATL were preserved after regressing out three other RDMs in the oral picture naming experiment. The overlap analysis further confirmed that the task-invariant neural representation of graph-CN RDM was confined in ATL, including the right TP and aMTG, as well as the left hippocampus.

##### Graph-shortest-path distance

In the oral picture naming experiment, RSA mappings between graph-SP RDM and neural RDMs revealed significant clusters in the pars triangularis of the left IFG, pMTG and pFG after regressing out three other RDMs. In the word familiarity judgment experiment, the significant clusters in multiple frontal-temporal regions were preserved, including medFG, OFC, SMA, IFG and pMTG, as well as widespread clusters located in the parietal cortex. The overlap analysis revealed the task invariant neural representation of graph-SP RDM in the pars triangularis part of the left IFG, left SMA and left pMTG/ITG.

##### Word2vec cosine distance

No clusters survived convention cluster extent-based inference threshold in either experiment.

In summary, the raw effects (across fMRI experiments) of the different language computation models were observed in both overlapping and different brain regions. The unique effects analyses revealed interesting dissociations: language graph-CN exhibited unique, task-invariant effects in the bilateral ATL, and language graph-SP exhibited unique effects in the left IFG and left pMTG/ITG (see Fig. 2c for bar plots of the unique effects). Edge (simple co-occurrence) and word2vec did not show overlapping regions of unique effects, i.e., those that cannot be explained by other models. Note that the left SMA showed the unique effects of language graph-SP but did not exhibit significant raw effects, which may result from complicated intercorrelations between these language computation RDMs, and it was not included in the following ROI analyses.

### Test of language specificity: Comparisons between language and visual computation models

To investigate whether the observed neural effects of the language computation models were specific to computing cumulative language-derived information or reflecting certain domain-general computations for cumulative information from any type of input, we constructed the same kind of graph representation using visual co-occurrence statistics from a large image corpus (a visual graph with 82,494 nodes and 3,920,082 visual co-occurrence edges, which derived from an image dataset – VisualGenome with 108,077 images) (Krishna et al., 2017). Same graph-related measures were also calculated, including visual edge, visual graph-CN and visual graph-SP.

The visual models were intermediately correlated with each other (visual edge RDM with visual CN RDM, Spearman *r* = 0.38; visual edge RDM with visual SP RDM, Spearman *r* = 0.46; visual CN RDM with visual SP RDM, Spearman *r* = 0.52; *Ps* < 2.48 x 10^-155^) and were weakly yet significantly correlated with the language models (visual edge RDM with language edge RDM, Spearman *r* = 0.15; visual CN RDM with language CN RDM, Spearman *r* = 0.23; visual SP RDM with language SP RDM, Spearman *r* = 0.12; *Ps* < 4.73 x 10^-16^).

In the whole-brain searchlight RSA of the visual RDMs (Fig. S4), the only significant cluster was the visual graph-CN in the right TOS (t_(25)_ = 4.96, peak MNI, x = 18, y = -99, z = 21, cluster size = 113 voxels) in the oral picture naming experiment under the convention cluster-extent-based inference threshold (voxelwise *p* < 0.001, FWE-corrected cluster-level *p* < 0.05).

We then focused on the ROIs from the language-model RSA unique effects (Fig. 2c): bilateral ATL (language graph-CN effect), left IFG and left pMTG/ITG (language graph-SP effect), to test whether they are also sensitive to visual graph-CN and visual graph-SP RDM. The ROI results (Fig. 3c) showed that none of these visual RDMs were significantly correlated with neural RDMs in the language overlapping regions (*Ps* > 0.096, uncorrected). Furthermore, the RSA results of the language models were preserved after regressing corresponding visual RDMs (*Ps* < 0.05, Bonferroni corrected).

**Fig. 3.**
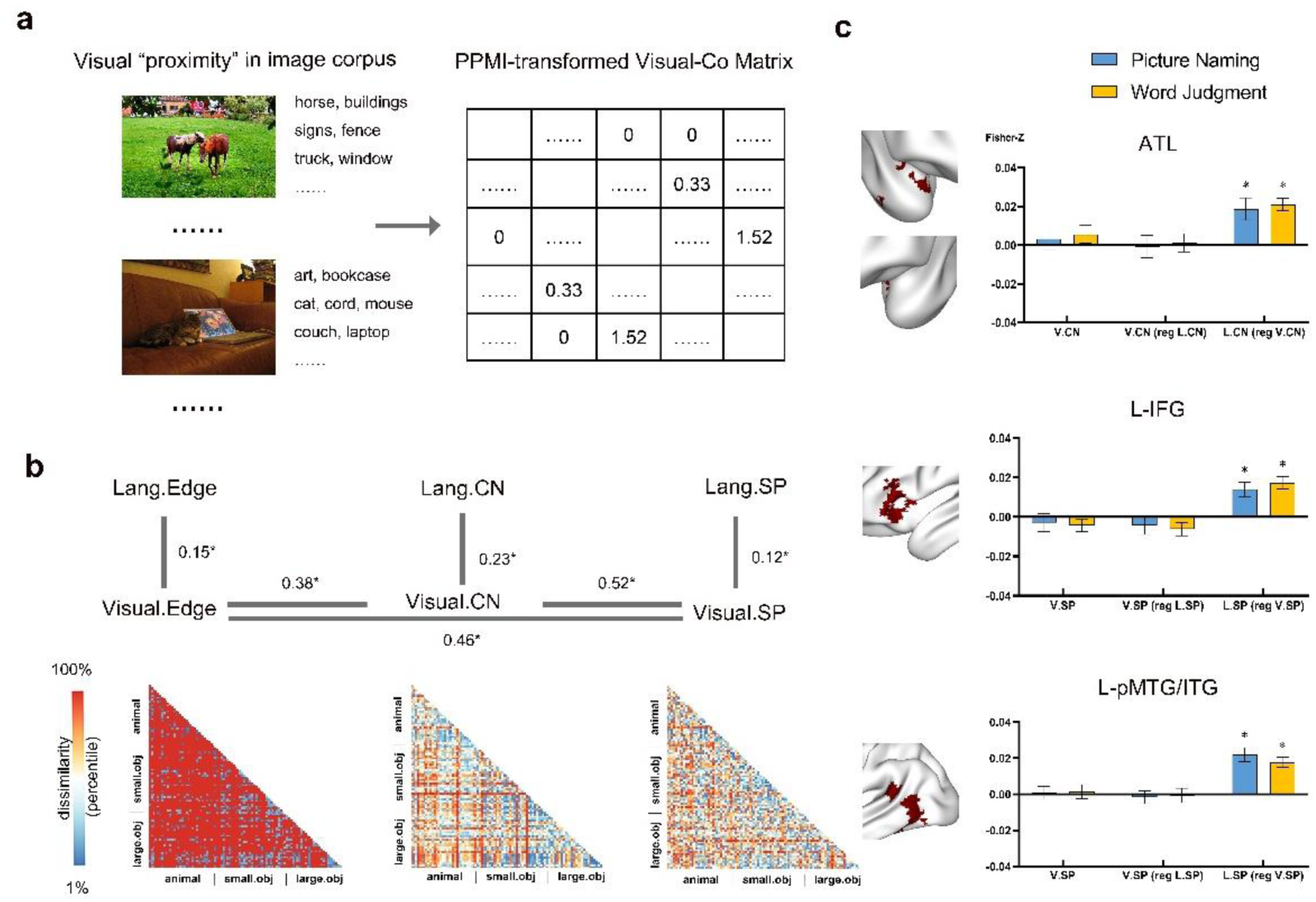
Neural effects of language computation models were not explained by visual co-occurrence statistics. (**a**) Visual co-occurrence statistics derived from daily experience. The raw co-occurrence counts in a given image “context” were calculated and normalized into PPMI values as our long-term coding of visual co-occurrence that is how likely two items co-occur visually. A visual co-occurrence matrix was calculated based on a human annotated image database -Visual Genome (approximately 108,077 images), resulting 82,494 objects and 3,920,082 non-zero visual co-occurrence relationship. (b) Results of visual RDMs and relationships with language RDMs. Like computation of language inputs, simple co-occurrence and graph-related measures were adopted to the visual statistics, including visual simple co-occurrence (edge), visual graph-CN and visual graph-SP). The RDMs were plotted and presented with the value of dissimilarity transformed into a percentile. Red indicates high dissimilarity (low similarity), and blue indicates low dissimilarity (high similarity). Correlations among these visual RDMs and language RDMs were investigated using Spearman rank correlations. (c) ROI results of visual RDMs in ROI regions from the language-model RSA unique effects. None of the neural effects of visual statistical models reached the significance level in the regions found to be sensitive to language graph-related distance (see Fig. 2c) (*p* > 0.05, uncorrected).

### Validation analysis

To test the robustness of the language graph representation, validation analyses were performed to address the following concerns: (1) Are the results affected by the graph type (weighted-graph vs binary-graph)? (2) Are the results affected by the specific window size choice in calculating the simple co-occurrence? (3) Are the results affected by the graph size and the method to select which words included in graph representation?

#### Effects of graph types

The language graph was built with weighted edges in the main results, and here, we reran the searchlight-based RSA mapping analysis using the binary versions of edges, graph-CN and graph-SP. Using ROI-based analysis, we found significant unique effects of graph-CN in the bilateral ATL and graph-SP in the left IFG, as well as the pMTG/ITG (Fig. 4a). We also conducted a whole brain searchlight analysis and the overlapping maps across two experiments were similar, except that the unique effects of SP were additionally found in the bilateral parietal cortices (Fig. S5).

**Fig. 4.**
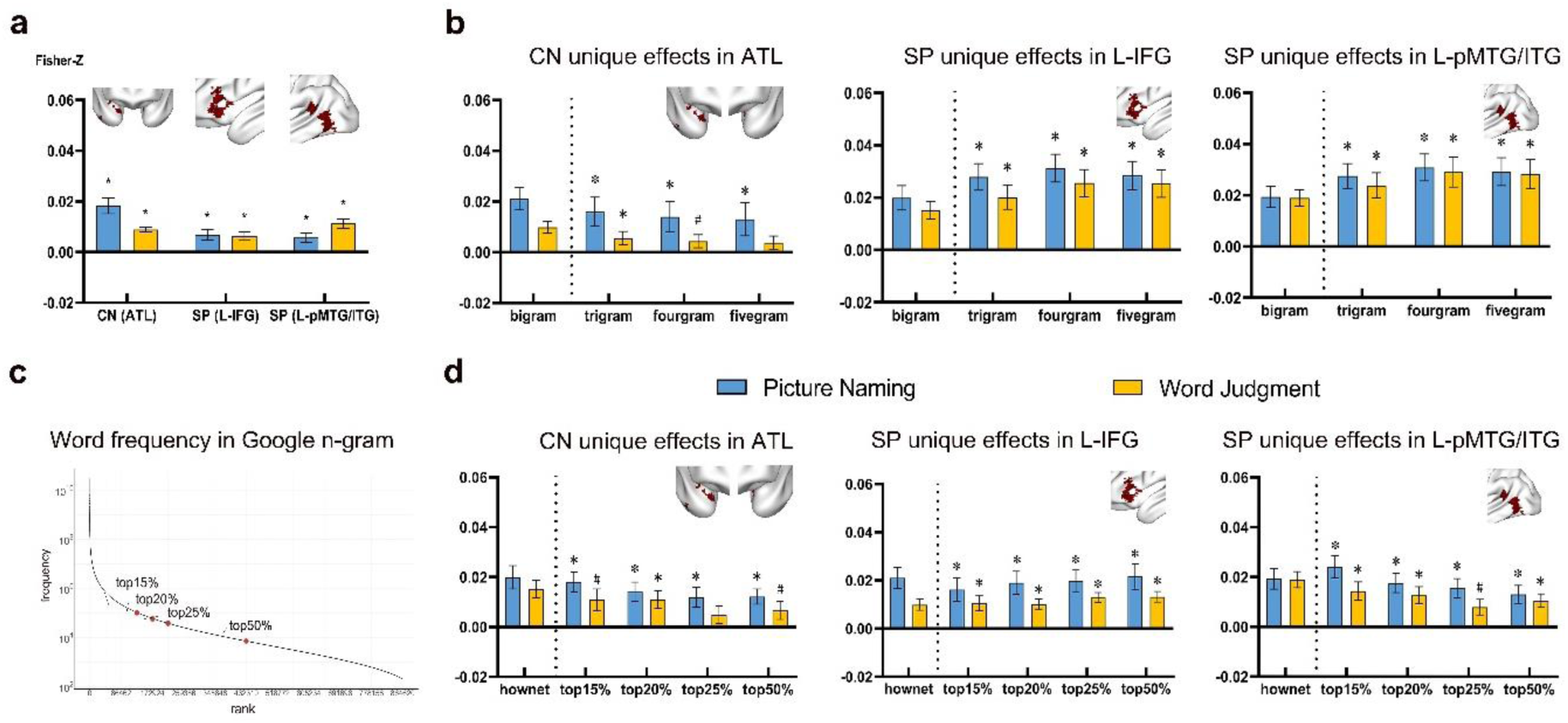
Validation analysis of graph representation using different graph types, window sizes, graph sizes and keyword selection methods. (**a**) ROI results using a different graph type (binary version). Significant RSA effects were found in the ROIs obtained in the main analyses (the language-model RSA unique effects) and the binary graph-CN and the binary graph-SP. (**b**) ROI results using different window sizes. Unique neural effects of the language-model RSA remained when the window size changed incrementally from two to five. (**c**) Distribution of word frequency in the Google n-gram corpus. Word frequency counts were log-transformed and only Chinese words were included, resulting a total of 864,620 words. We selected keywords based on word frequency and changed graph size incrementally (top15%, top20%, top25%, top50%) to validate the effects of word selection bias and the robustness under different graph sizes. (**d**) ROI results using graphs of different sizes. Different graph sizes yielded similar results in the language ROIs. (*, *p* < 0.05, Bonferroni corrected; #, *p* < 0.05, uncorrected). The bar plots for the original bigram word co-occurrence which intersected with OpenHowNet corpus are presented on the left side of the dashed line for visualization without additional statistical inference.

#### Effects of window sizes

The main analyses considered immediate co-occurrences (window size = 2). Here, we reconstructed language graphs based on trigram, fourgram and fivegram word co-occurrence patterns and reran the searchlight-based RSA mapping analyses in both fMRI experiments. The main patterns of ROI-based results were largely maintained across all window sizes: significant unique effects of graph-CN were found in the bilateral ATL when the window sizes were 2-4, with the effects decreasing when the window sizes changed to 5; significant unique effects of graph-SP were found in the left IFG and left pMTG/ITG (Fig. 4b). Overlap results of whole brain searchlight RSA using different window sizes also yielded comparable results to those in the main analysis.

#### Effects of graph sizes and word selection methods

In the main analyses, we selected the word in the Google n-gram corpus based on the intersection part of a human annotated Chinese Knowledge Database (OpenHowNet, including 223,767 unique Chinese and English words) (Qi et al., 2019). 83,007 Chinese words were used to construct the language graph to reduce the computational loads. Here, we adopted a different keyword selection method (select words based on word frequency) and calculated the co-occurrence counts using the top 15%, 20%, 25% and 50% most frequent words from the total of 864,629 Chinese word samples. The log transformed frequency was plotted and visualized in Fig. 4c. We then reconstructed our graph based on these different frequency ranges. Again, the unique effects of graph-CN in the bilateral ATL and the unique effects of graph-SP in the left IFG and left pMTG/ITG for SP remained significant across all graph sizes (Fig. 4d). Overlap results of whole brain searchlight RSA under these different graph sizes were similar to those results reported in the main analysis.

#### Effects of corpus selection

In the main analysis, we adopted a published, open accessed word2vec model which was pre-trained using a language corpus different from the one from which the other two models derived. It is possible that the different sizes and text contents between two corpuses contribute to the different neural results between graph models and vector-embedding models. To eliminate this possibility and test the robustness of vector-embedding representation, we trained the word2vec model based on the same Google n-gram corpus (see Fig. S6 for detailed training parameters). Both ROI-based RSA results and whole brain searchlight RSA results were largely replicated (Fig. S6).

## Discussion

To understand whether and how the human brain represents word meanings derived from cumulative language inputs, we mainly tested three different types of computations that extract language statistical patterns from a large corpus in fitting word-processing brain activity patterns: simple co-occurrence, two graph-space relations (graph-common-neighbors and graph-shortest-path), and neural-network-vector-space relations. In two fMRI experiments, oral picture naming and written word familiarity judgment, which vary by input and output peripheral processes (visual picture recognition, phonological lexical access and output; visual word recognition, button press) but share common word meaning representation, we observed that word relations constructed from all four models correlated words’ brain activity patterns across broadly distributed brain regions. However, the word relations derived from a topological raw graph space, and not the other two types, have unique explanatory power for the neural activity patterns in brain regions that have been shown to be particularly sensitive to language processes, including ATL, IFG, and pMTG/ITG. Intriguingly, the organization of different graph relations was respected by these regions with ATL based on the proportions of common neighbors in a graph and with IFG and pMTG/ITG based on the shortest path distances. These neural results of language graph representation were robust across different language co-occurrence measuring window sizes and graph sizes and were relatively specific to language inputs, as they were not associated with relational structures derived from visual co-occurrence statistics when using the same computation methods.

Our results describing correlations between all four language model patterns and broadly distributed brain regions are consistent with the literature findings (Anderson et al., 2019; Carota et al., 2017; Carota et al., 2021; Huth et al., 2016; Pereira et al., 2018; Xiaosha Wang et al., 2018). These studies used one or two types of language computation models. It is not clear what specific kinds of computations are driving these effects, given the medium-to-high correlations among different types of language statistical models. Our study, by contrast, compared the effects of three different kinds of computations of the language corpus and revealed that graph-related measures, constructed from simple co-occurrences, specifically capture the neural activity patterns in brain regions that are related to language - the effects of simple co-occurrence and w2v could be explained by graph relations but not vice versa. Furthermore, different graph relations are captured by neural activities in different brain regions: the neural activity similarity between two words in ATL is predicted by the proportion of common neighbors these two words have in the word co-occurrence graph; in IFG and pMTG/ITG, the similarity is predicted by the distance of the shortest path between two words in the graph (see below).

What do the two graph relations reflect, how are they different from word2vec cosine distances, and what are the implications for the representations in these three brain regions? The graph representation retained the original dimensions given the size of the word co-occurrence matrix (Jackson & Bolger, 2014), with the extraction of statistical information achieved through relationships between highly informative neighbors and paths. In this way, more “historical information” can be preserved. Furthermore, we can select subgraphs or pruning the edges based on word frequency without dramatically changing the network structures, especially the interconnected neighborhoods. By comparison, word embedding techniques project the word co-occurrence statistics into a dense vector space, resulting a holistic representation of word meanings. In the case of word2vec, statistical information was compressed into a fixed, arbitrary, usually 300-dimension vector space through error-driven training and hyperparameter optimizations. While some studies suggested that word2vec models, especially skip-gram versions, mainly capture the second-order co-occurrence (Schlechtweg, Oguz, & Walde, 2019), our between-model correlation results found both first-order and second-order measures (i.e. common neighbors) were significant correlated with word2vec models. It remains controversial whether these hyperparameter tuning processes are psychological meaningful and what information is retained and lost after dimension reductions (Kumar, 2021).

Graph computation also has the advantage of unpacking different types of topological computations, which provides novel computational insights into the functionality of ATL, IFG and pMTG. These three regions are among those consistently activated by language processing in general (Fedorenko, Hsieh, Nieto-Castañón, Whitfield-Gabrieli, & Kanwisher, 2010) and, in particular, more strongly activated by abstract words than concrete words (Binder et al., 2009; Hoffman et al., 2015; J. Wang et al., 2010). While these activations have been speculated to be related to greater linguistic reliance on abstract words, the detailed functional roles remain controversial. Here, we observed that while they are all sensitive to language computation models, ATL represents words based on their proportions of shared common neighbors, and IFG & pMTG/pITG represents words based on their shortest path. Considering cat (‘猫’), mouse (‘老鼠’) and mousetrap (‘夹子’), mouse-cat have more common neighbors than mouse-mousetrap, whereas mouse-mousetrap has a greater distance of the shortest path than mouse-cat (i.e., it takes more steps to go from mouse to cat than to mousetrap). Thus, brain activity when accessing mice is more similar to that of cats in ATL and to that of mice and mousetrap in IFG and pMTG/ITG. The shortest path captures long-range dependency in graph space - it measures association strength between two non-adjacent nodes; the common neighbors capture structural similarity based on second-order proximity. These two types of statistical properties have been shown to be associated with different types of meaning relations, with CN with taxonomy/semantic categorical similarity and edge with associative relations (Jackson & Bolger, 2014). Here, we also performed an ad hoc analysis on the word sets with rated taxonomical/thematic relations (Xu et al., 2018) and found that CN was correlated with taxonomic relations (Spearman *r* = 0.48, *P* < 0.001), weakly correlated with thematic relations (Spearman *r* = 0.10, *P* = 0.003) after regressing out SP measure, and SP was positively correlated with thematic relations (Spearman *r* = 0.41, *P* < 0.001), not taxonomical relations (Spearman *r* = -0.08, *P* = 0.98) after regressing out CN measure. These observations align with the findings that word (semantic, categorical) similarity was computed in ATL (C. B. Martin, Douglas, Newsome, Man, & Barense, 2018; Xu et al., 2018) and that IFG and/or pMTG/ITG contribute to the retrieval of infrequent word associations, i.e., longer path length distance (Badre, Poldrack, Paré-Blagoev, Insler, & Wagner, 2005; Whitney, Kirk, O’Sullivan, Lambon Ralph, & Jefferies, 2011). Finally, these findings are consistent with the topological structural observations of the intrinsic functional semantic network, in which these regions were identified as connector hubs that bind together different brain subnetworks: ATL binds the perisylvian language network and multimodal experiential network, and pMTG binds the perisylvian language network and frontoparietal control network (Xu, He, & Bi, 2017; Xu, Lin, Han, He, & Bi, 2016). The current results move beyond these descriptive notions contrasting taxonomy vs. associative and representational vs. control and, for the first time, provide a mathematical computational account for how they represent information derived from cumulative language experience.

Are these brain regions sensitive to graph-topological structures from all stimuli domains? On the one hand, several sequential learning experiments have shown that changes in neural representation in medial temporal, anterior temporal, and frontal regions, after a short training session involving sequence exposure (Schapiro, Kustner, & Turk-Browne, 2012; Schapiro et al., 2013), follow graph-based statistical regularities (simple edge, community structure) of arbitrary visual stimuli sequences during the formation of episodic memory (see also (Garvert et al., 2017), where MTL was sensitive to the path distance of daily object sequences). On the other hand, we did not observe any sensitivity to common neighbors or shortest path measures from the large visual co-occurrence database. It is possible that these regions are indeed only sensitive to the language graph structures and that the results observed in the studies above are driven by the potential language encoding applied by the subjects in those experiments. More generally, a wave of recent studies has highlighted the exoplanetary power of “spatial relationship” or “grid-like” structures in representing conceptual knowledge and information in memory in general (Constantinescu, O’Reilly, & Behrens, 2016; Theves, Fernandez, & Doeller, 2019; Theves, Fernández, & Doeller, 2020). Here, we showed that the topological distances in a graph space are actually better predictors than the cosine distance in the vector embedding space in explaining word representations. The breadth of the application of the observed computational structures in representing information in these regions warrants further testing (Peer, Brunec, Newcombe, & Epstein, 2020).

A few methodological caveats need to be considered. First, in our current investigation, we used a large-scale language corpus as the proximity of collective language experience on a group level. It may not be an accurate reflection of specific language inputs at the individual level. Computational modeling of word meaning representations in the future could benefit from collecting and estimating individual participants’ language inputs with the help of personalized big data techniques. Second, we specifically considered relatively simple models that are fully data driven, without prior knowledge such as grammatical information and attentional allocation mechanisms. In recent years, there has been a surge of computational models with improved performances in various language tasks, such as recurrent neural-network models (ELMo) (Peters et al., 2018) and attention neural-network models (BERT) (Devlin, Chang, Lee, & Toutanova, 2018), and whether their computational architecture is relevant to brain computations in ATL/IFG/pMTG awaits further study.

To conclude, across two fMRI experiments investigating word meaning access, we tested the potential computational mechanisms through which cumulative language experience is captured by the human brain, evaluating the effects of three major types of statistical pattern extracted in the large-scale language corpus. Graph-based topological models had unique explanatory power on words’ neural activity patterns beyond simple co-occurrence and vector-embedding models, showing effects in the anterior temporal lobe (capturing graph-common-neighbors), inferior frontal gyrus and posterior middle/inferior temporal gyrus (capturing graph-shortest-path). The latter two did not show effects beyond graph models. These results highlight the role of cumulative language inputs in shaping word meaning representations in this set of brain regions and provide a computational account of how they capture different types of language-derived information.

## Materials and Methods

### Participants

Twenty-nine participants (19 female; median age, 20 years; range, 18-32 years) were recruited in our study and were scanned in 2 fMRI experiments on separate days. All participants were right-handed native Mandarin Chinese speakers with normal or corrected-to-normal vision and had no history of neurological or language disorders. They provided written informed consent. This study was approved by the institutional review board of the State Key Laboratory of Cognitive Neuroscience and Learning, Beijing Normal University (ICBIR_A_0040_008), adhering to the Declaration of Helsinki for research involving human subjects. Three participants in the oral picture naming experiment and 3 participants in the word familiarity judgment experiment were excluded from data analysis due to successive head motions (> 3 mm/3°). Another 6 participants in the word familiarity judgment experiment were excluded because of poor behavioral responses (> 20% nonresponsive trials in more than one run). Therefore, the fMRI data of 26 participants in the oral picture naming experiment and 20 participants in the word familiarity experiment were analyzed.

### Stimuli

Ninety-five objects were selected in our fMRI experiments, including 3 common categories (32 animals, 35 small manipulable artifacts and 28 large nonmanipulable artifacts). In two separate fMRI experiments, objects were shown as words and colored pictures (see Fig. S1 and Table S1 for details). Pictures were 400 x 400 pixel images with the representative exemplar of the object presented against a white background (10.55° x 10.55° of visual angle). Words were presented in white FANG SONG font against a black background and subtended approximately 7.92° x 2.64° of the visual angle.

### Computation of language models

Three types of language computation mechanisms were adopted to extract four kinds of statistical patterns between words: simple word co-occurrence, graph-related (graph-common-neighbors and graph-shortest-path) space and word2vec cosine vector space. Distances between 95 object pairs were derived from these four measures to construct the theoretical RDMs for the subsequent RSA computation with the neural data.

#### Construction of the PPMI-normalized simple co-occurrence matrix

Raw word co-occurrence counts were first collected in the Chinese Web Google n-gram corpus (https://catalog.ldc.upenn.edu/LDC2010T06) (Liu et al., 2010). The corpus included publicly accessible documents (a total of 882,996,532,572 tokens, 1,616,150 unique tokens including 864,629 unique Chinese words) on the Chinese Internet by the end of 2008. In this dataset, n-grams within a context window ranging from 1 to 5 were extracted by the original developer using an auto segment parser. Bigram word co-occurrence counts were used in the main results. We also calculated our measures based on trigram, fourgram and fivegram word co-occurrence counts in the validation analysis. To reduce the computational loads and filter low-frequency and meaningless words, we further selected the keywords based on a human annotated Chinese Knowledge Database (OpenHowNet, 223,767 unique Chinese and English words, (Qi et al., 2019)), resulting in 83,007 Chinese words and a downsampled corpus with a size of approximately 587 billion words. In the validation analysis, we also adopted different keyword selection methods based on different word frequency ranges and calculated the co-occurrence counts using the top 15%, 20%, 25% and 50% most frequent words of a total of 864,629 Chinese word samples.

Importantly, pointwise mutual information (PMI)-normalized word co-occurrence counts between two nodes, u and v, were adopted to construct the 83,007 x 83,007 simple co-occurrence matrix, which reflects the direct proximity between two words in long-term language exposure (Church & Hanks, 1990). Here, we only consider positive PMI (PPMI) to obtain the sparse representation of the word co-occurrence matrix, as it is more computationally efficient. Additionally, the negative values might introduce “uninformative” noise (Levy & Goldberg, 2014).

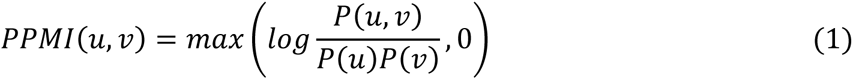

#### Construction of the graph representation

To calculate graph-related distance, 83,007 words in the word co-occurrence PPMI matrix were taken as nodes, and 34,586,840 word-pair PPMI values were taken as edges to construct the word graph space. Considering that between-word relations in the graph space are remarkably rich, we mainly considered two graph-related measures based on current knowledge and algorithms from graph theory and semantic network practice (Jackson & Bolger, 2014; Liben-Nowell & Kleinberg, 2007; Lü & Zhou, 2011; Newman, 2001): The Jaccard similarity coefficient of common neighbors (graph-common-neighbors) and the shortest path distance (graph-shortest-path). For a given word pair, the Jaccard similarity coefficient of common neighbors was calculated as the summed PPMI weights of their shared edges of neighbors divided by the summed PPMI weights of unions edges of neighbors connected with two words, which reflects the second-order proximity between the two words (Jackson & Bolger, 2014). Precisely:

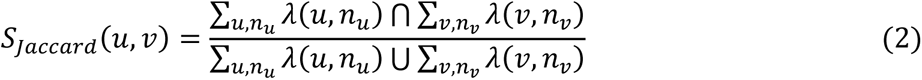

where *n_u_* is the neighbor node of *u*, and *λ(u, n_u_)* is the weighted PPMI value (edge) between *u* and *n_u_*. The same is for *n_v_* and *λ(u, n_v_)*.

For the calculation of weighted shortest path distance (Newman, 2001), we summed weights along the shortest path between two words in a graph-defined space with inverted PPMI using Dijkstra’s algorithm in Neo4j (http://neo4j.com/). The measure was precisely calculated as:

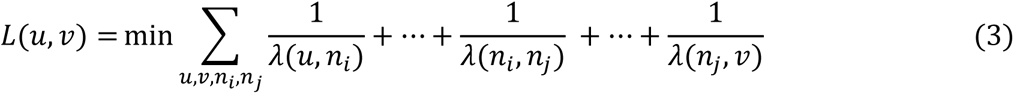

where *λ(u, n_i_)* is the weighted PPMI values (edge) between *u* and a given node *n_i._*.

Notably, both first-order edges and common neighbors measure the similarity between two words, while the weighted length of shortest paths measures the distance between two words. In the following analysis, we converted the former two into dissimilarity representations using a 1 minus calculation to obtain the corresponding RDMs.

Another proximity measure between words is Katzβ (communicability), which is calculated from ensembles of all paths with a damping factor β to discount the path weights. We did not consider this measure for the following reasons: 1) it is computationally expensive to calculate the global sums over the collection of all paths given the high degree of interconnection in the language graph; 2) the selection of dumping factor β is usually arbitrary and unexplored; and 3) the measure is similar to common neighbors when β is very small (Liben-Nowell & Kleinberg, 2007).

Note that the graph-related measures we adopted here were based on the weighted version of graph space (i.e., the edge in the graph was a PPMI-normalized word co-occurrence value) to preserve as much statistical information as possible. In the validation analysis, we also calculated graph-based measures with the binary version (the edges with PPMI greater than 0 were coded as 1 and other edges as 0).

#### Construction of the vector-embedding (word2vec) representation

Another way to extract language statistical information is to project the word co-occurrence statistics in a hidden layer space using word embedding techniques. In this study, we adopted a pretrained, open-accessed word2vec vector dataset that achieved state-of-art performance on analogical reasoning of Chinese semantic relations (Li et al., 2018). A large mixed corpus (4037 million words) was selected from Baidu Encyclopedia, Chinese Wikipedia, People’s Daily News, Sogou News, Financial News, Zhihu_QA, Weibo, and 8599 modern Chinese literature books. The basic parameter settings of this model were as follows: dynamic window size = 5, subsampling rate = 10^-5^, negative sample number = 5, iteration = 5, low-frequency word = 10, dimension = 300). The skip-gram architecture (Mikolov et al., 2013) was adopted to predict the surrounding context words given an input target word. After intensive training and prediction, each word was represented as a 300-dimensional vector, and word relations were calculated as the cosine distance of these vectors. To confirm our neural findings with the word2vec model and make results between word2vec and the other three measures more comparable, we further repeated our analyses on a word2vec model trained on the same corpus (see Fig. S6 for details).

### Computation of visual models

To examine whether our findings of language computation models were specific to language inputs or reflect domain-general computations for any kind of statistical co-occurrence pattern, we considered another kind of co-occurrence (visual co-occurrence statistics). Visual co-occurrence was collected based on an image database - VisualGenome (http://visualgenome.org/) (Krishna et al., 2017). The image database consisted of 108,077 images; objects in each image were annotated by human observers, and the labels were further mapped to Wordnet synsets. We first extracted the unique labels across the database (82,494 objects in total). The co-occurrence counts between objects were first obtained for each image and then summed over all images to obtain the raw object co-occurrence matrix. Similar to the language graph representation, we further constructed visual models using the same measures: visual edge (PPMI value, yielding 3,920,082 visual co-occurrence relationships), visual graph-common-neighbors and visual graph-shortest-path. RDMs were constructed accordingly.

### Low-level control RDMs

We constructed the following low-level control RDMs (Fig. S1). First, to control for the low-level visual similarity effects between pictures/words, we calculated the pixel and gist dissimilarity of image pairs and the pixel dissimilarity of word pairs separately. The pixel dissimilarity was computed by the Pearson correlation distance between the grayscale values of images. The gist dissimilarity was computed by the Euclidean distance of 32 visual features of images (4 kinds of frequency, 8 kinds of orientation) (Oliva & Torralba, 2001). Additionally, word familiarity judgment responses collected during fMRI scanning were used to construct the button-press RDM based on the absolute difference between the group-averaged familiarity scores of each object pair.

### fMRI experimental design

We adopted a condition-rich event-related fMRI experimental design to estimate the hemodynamic responses for each item (Kriegeskorte, Mur, & Bandettini, 2008). Two experiments were carried out: an oral picture naming experiment and a word familiarity judgment experiment. In the oral picture naming experiment, participants were instructed to overtly name the objects in colored pictures as precisely and quickly as possible. In the word familiarity judgment task, participants were asked to judge whether the presented written name of objects was familiar according to their personal experience by pressing the corresponding buttons (familiar: left index finger; unfamiliar: left middle finger).

The two experiments were conducted separately on two days for each participant, and the word familiarity judgment experiment was always carried out before the oral picture naming experiment to avoid eliciting the visual imagery of the object in the word experiment. Each experiment had 6 runs, with each word/picture repeated 6 times. Each run (8 min 48 s) consisted of 95 trials, and each word/picture was presented exactly once. In each trial, there was 0.5-s fixation and 0.8-s stimulus presentation, followed by the intertrial interval (ITI) ranging from 2.7 to 4.7 s. For the order of stimuli and length of ITI, we first determined the sequence of the 3 categories using the optseq2 optimization algorithm (http://surfer.nmr.mgh.harvard.edu/optseq/) (Dale, 1999) and further randomized the orders of items in each category. Each run began and ended with a 10-s fixation period. The presentation and timing of stimuli was implemented using E-prime 2 (Psychology Software Tools) (Schneider, Eschman, & Zuccolotto, 2002).

### Image acquisition

Functional and structural MRI images of two experiments were collected for each participant using a 3T Siemens Trio Tim Scanner at the Beijing Normal University MRI Center. A high-resolution 3D structural image was collected with a 3D-MPRAGE sequence in the sagittal plane (144 slices, TR = 2530 ms, TE = 3.39 ms, flip angle = 7°, matrix size = 256 x 256, voxel size = 1.33 x 1 x 1.33 mm). Functional images were acquired with an echo-planar imaging (EPI) sequence (33 axial slices, TR = 2000 ms, TE = 30 ms, flip angle = 90°, matrix size = 64 x 64, voxel size = 3 x 3 x 3.5 mm with gap of 0.7 mm).

### Image data analysis

#### Preprocessing

Task-fMRI data were preprocessed and analyzed for each experiment using Statistical Parametric Mapping (SPM12; http://www.fil.ion.ucl.ac.uk/spm). For each individual participant, the first 5 volumes (10 s) of each run were discarded for signal equilibrium. The preprocessing of functional images included slice timing correction and head motion correction, and the resulting unnormalized and unsmoothed images were entered into general linear models (GLMs). The structural images were segmented into different tissue types; the resulting gray matter probabilistic images were coregistered to the mean functional image in the native space, resliced to the spatial resolution of functional images, and thresholded at one-third to obtain the gray mask of each subject. The forward and inverse deformation fields of each subject’s native space to the Montreal Neurological Institute (MNI) space were also obtained at this step.

#### GLM

For the functional images in the native space in each subject, GLM was built to obtain object-level neural activation patterns. The GLM contained onset regressors for each of 95 items, 6 regressors of no interest corresponding to the 6 motion parameters, and a constant regressor for each run. Each object regressor was convolved with a canonical HRF, and a high-pass filter cutoff was set as 128 s. Additionally, to ensure the maximal coverage of regions with a low ratio of signal-to-noise (e.g., ATL), the SPM implicit mask threshold was set to 10% of the mean of the global signal (compared with the default threshold of 80%) (Devereux et al., 2013). For each experiment, the t-value images (each condition relative to baseline) were obtained to capture the neural activation patterns.

#### RSA searchlight analysis

To identify the brain regions that may represent different language computation models, we carried out RSA using a searchlight procedure (Kriegeskorte et al., 2006; Kriegeskorte, Mur, & Bandettini, 2008). For each voxel in the gray matter mask in the native space, the t-values of 95 objects within a sphere (radium = 10 mm) centered at that voxel were extracted and correlated across object pairs to create the 95 x 95 neural representational dissimilarity matrix using Pearson correlation distance. The neural RDM was then compared with language/image theoretical RDMs using partial Spearman rank correlation controlling for low-level control RDMs (raw effects) and for other language statistical models and low-level control RDMs (unique effects). The resulting *r*-values were assigned to the center voxel of the sphere, and the searchlight procedure across each gray matter voxel produced a gray matter *r*-map for each participant. These individual *r*-maps were Fisher-z transformed, normalized into the MNI space, and spatially smoothed using a 6 mm full-width half-maximum Gaussian kernel.

For group-level random-effects analysis, one-sample *t* tests were performed across the individual *r*-maps using permutation-based statistical nonparametric mapping (SnPM13; https://go.warwick.ac.uk/tenichols/snpm). No variance smoothing was used, and 10,000 permutations were performed. To localize the effects of theoretical models in each task, the RSA maps were thresholded at a conventional cluster extent-based inference threshold (voxelwise *p* < .001, FWE corrected cluster-level *p* < .05). To demonstrate the task-invariant effects of language computation models, the RSA maps of each fMRI experiment were thresholded at voxelwise *p* < .005, cluster sizes > 20 voxels, and overlapped with each other. The overlapping regions were considered to show significant positive correlations between neural and theoretical RDMs in both experiments. The brain results were projected onto the MNI brain surface using BrainNet Viewer (https://www.nitrc.org/projects/bnv) (Xia, Wang, & He, 2013).

#### RSA ROI analysis

For the regions showing the task-invariant effects (i.e. overlapping region) of language computation models identified by the RSA searchlight analysis, we carried out several region of interest (ROI) analyses, including the effects of the visual models and validation analyses with different graph types, window sizes, graph sizes and keyword selection methods. For each of these RDMs, we first performed the RSA searchlight mapping procedures to obtain the whole-brain *r*-maps in the native space for each subject. These *r*-maps were then Fisher-z transformed and normalized into MNI space. The Fisher-transformed correlation values were averaged across voxels within the ROI for each subject. One-sample *t* tests across subjects were then conducted to test whether the RSA results of the theoretical models were significantly above zero.

## Acknowledgments

We thank Xing Wang for training word2vec models in the validation analyses. This work was supported by the National Natural Science Foundation of China (31925020, 31671128 to Y.B., 31700943 to X.S.W., 32071050 to X.Y.W, 31700999 to T.W.), Changjiang Scholar Professorship Award (T2016031 to Y.B.), the National Program for Special Support of Top-Notch Young Professionals (to Y.B.), the Interdisciplinary Research Funds of Beijing Normal University (to Y.B.), and China Postdoctoral Science Foundation (2017M610791 to X.S.W., 2020M670190 to H.Y.). The funders had no role in the conceptualization, design, data collection, analysis, decision to publish or preparation of the manuscript.

## Author contributions

YB conceived and designed the study; ZF performed the study; XSW, XYW, and HY contributed to critical discussions; JW computed the visual relation measures; TW acquired fMRI data; XL contributed to graph measure computation; ZL and HC contributed to natural language computation modelling; YB and ZF wrote the paper.

## Additional information

Supplementary material (Fig S1 – S6; Table S1 – S3).

## Supplementary Materials

**Fig. S1.**
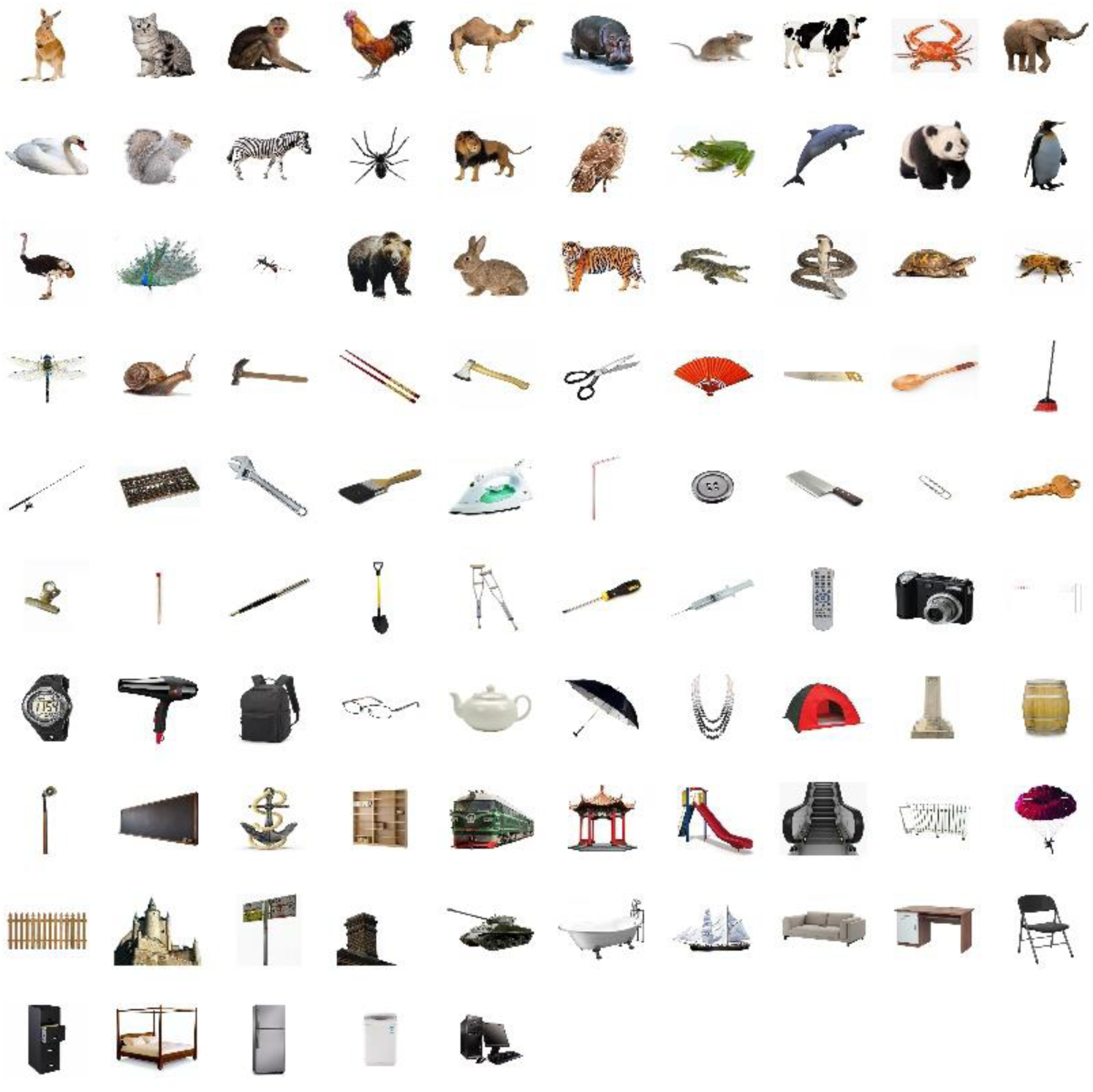
Picture stimuli presented in the oral picture naming experiment. Ninety-five object pictures were selected in the oral picture naming experiment, including 3 common categories (32 animals, 35 small manipulable artifacts and 28 large nonmanipulable artifacts). See Table S1 for the corresponding names of these pictures.

**Fig. S2.**
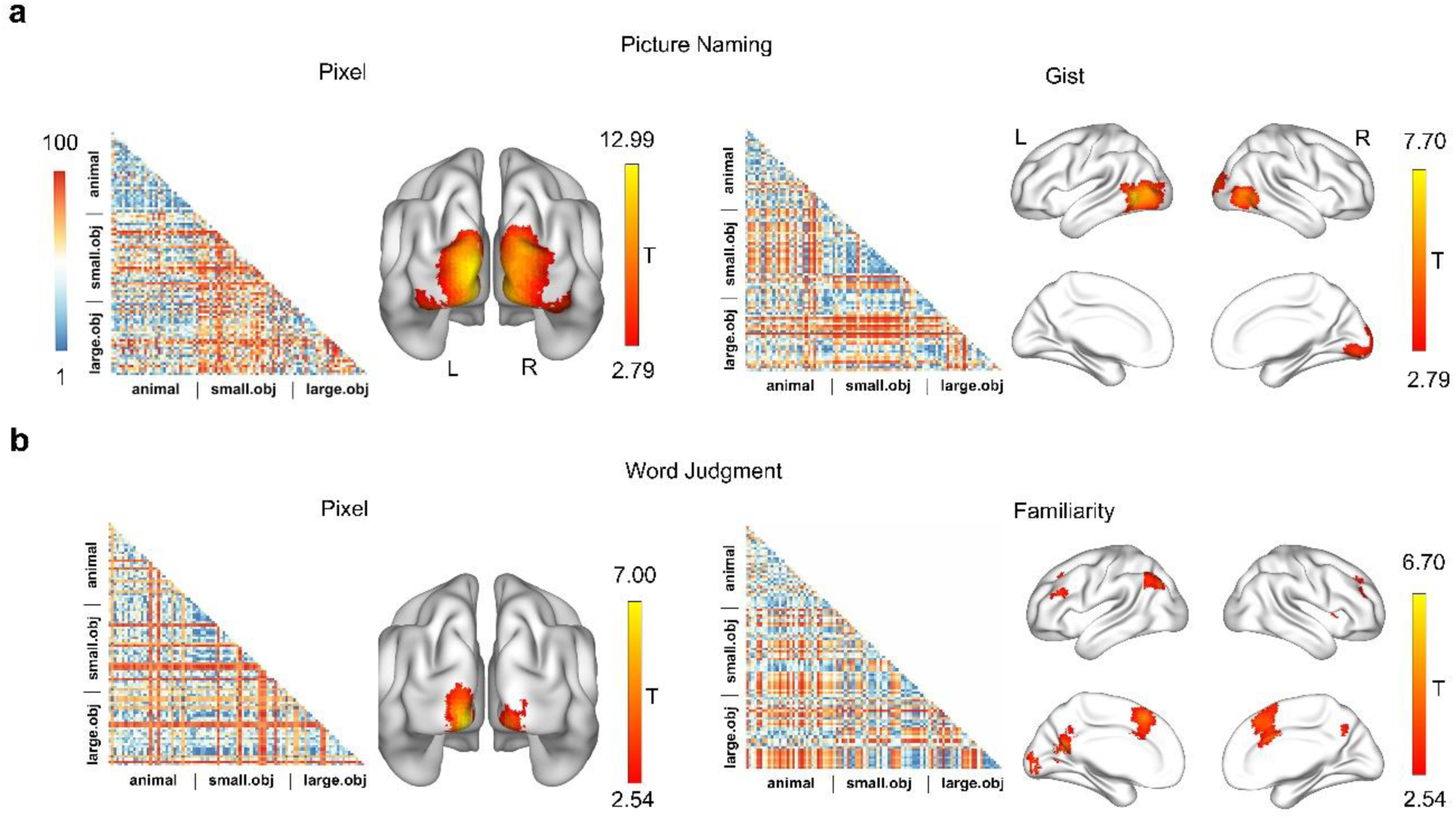
Low-level control models and their neural effects across two experiments. (**a**) Visualization of control RDMs in the oral picture naming experiment. The pixel and gist RDM were used to control the effects of local visual statistical information in the presented pictures. The value of dissimilarity was transformed to a percentile for display. Red indicates high dissimilarity (low similarity), and blue indicates low dissimilarity (high similarity). The neural effects of each RDM were presented (uncorrected voxelwise *p* < 0.005, one-tailed, cluster size > 20 voxels, for display purposes). (**b**) Visualization of control RDMs in the word familiarity judgment experiment. The pixel RDM was used to control the effects of local statistical visual information in presented words. The familiarity (button press) RDM was used to control the task-induced processing.

**Fig. S3.**
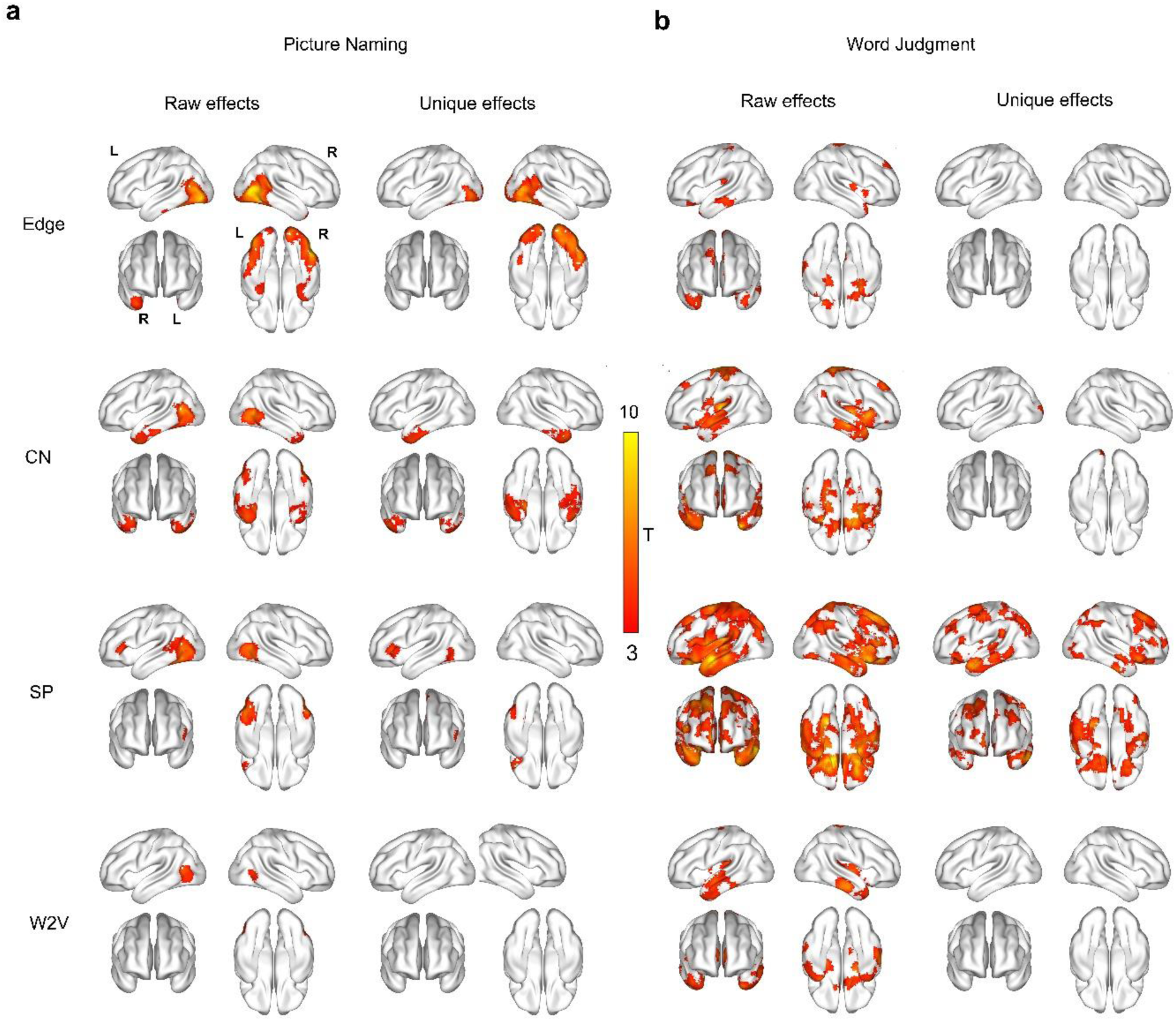
Whole brain searchlight RSA results of language computation models in each experiment. (**a**) The ‘raw effects’ and ‘unique effects’ (regressing the other three) of neural representation of different language model RDMs in the oral picture naming experiment. The convention cluster-extent-based inference threshold was adopted (voxelwise *p* < 0.001, FWE-corrected cluster-level *p* < 0.05). (**b**) The ‘raw effects’ and ‘unique effects’ of different language model RDMs in the word familiarity judgment experiment.

**Fig. S4.**
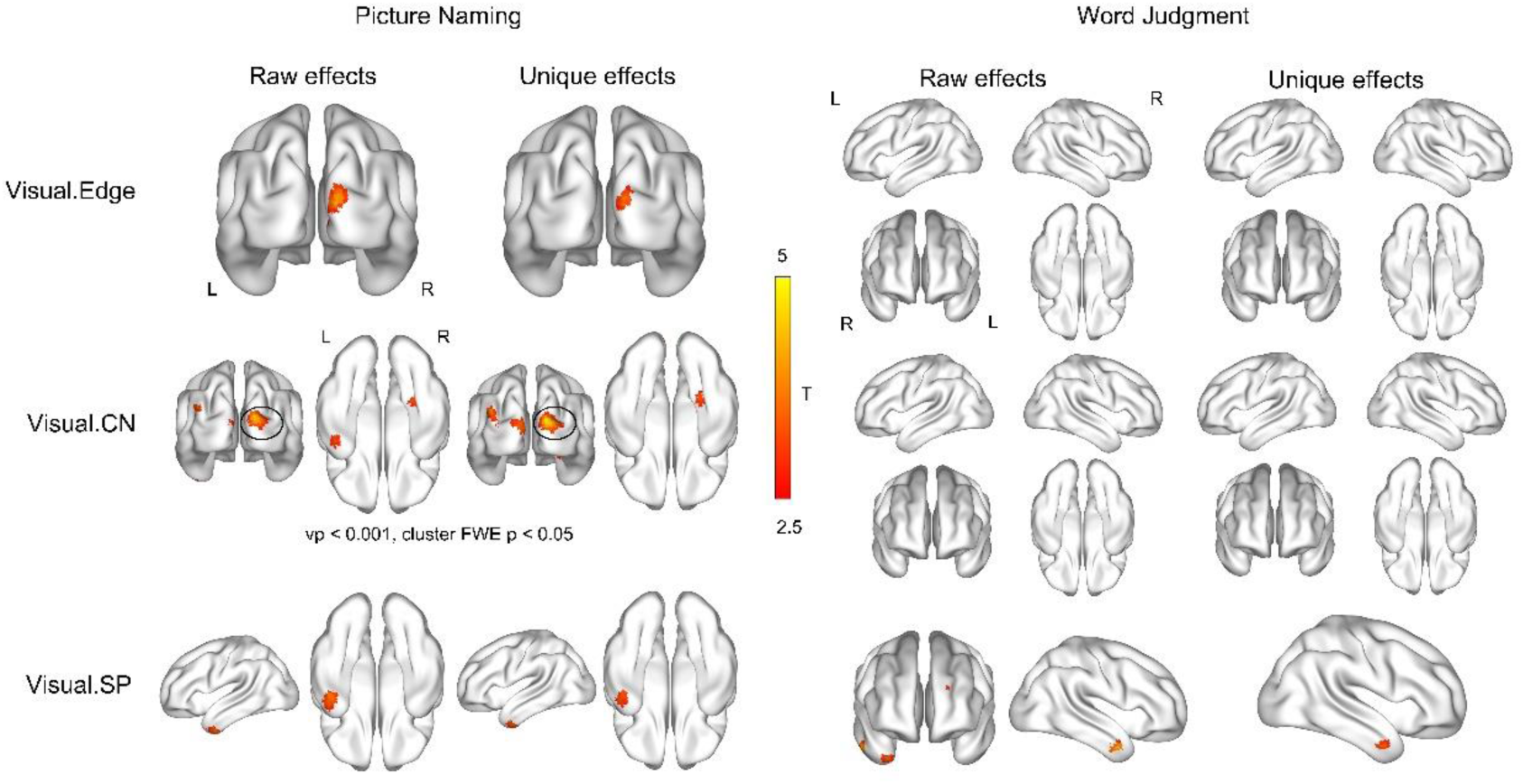
Whole brain searchlight RSA results of visual models in each experiment. The ‘raw effects’ and ‘unique effects’ (regressing the other two) of the visual models (visual edge, visual graph-CN and visual graph-SP) in two experiments are shown here (uncorrected voxelwise *p* < 0.005, one-tailed, cluster size > 20 voxels, for display purposes).

**Fig. S5.**
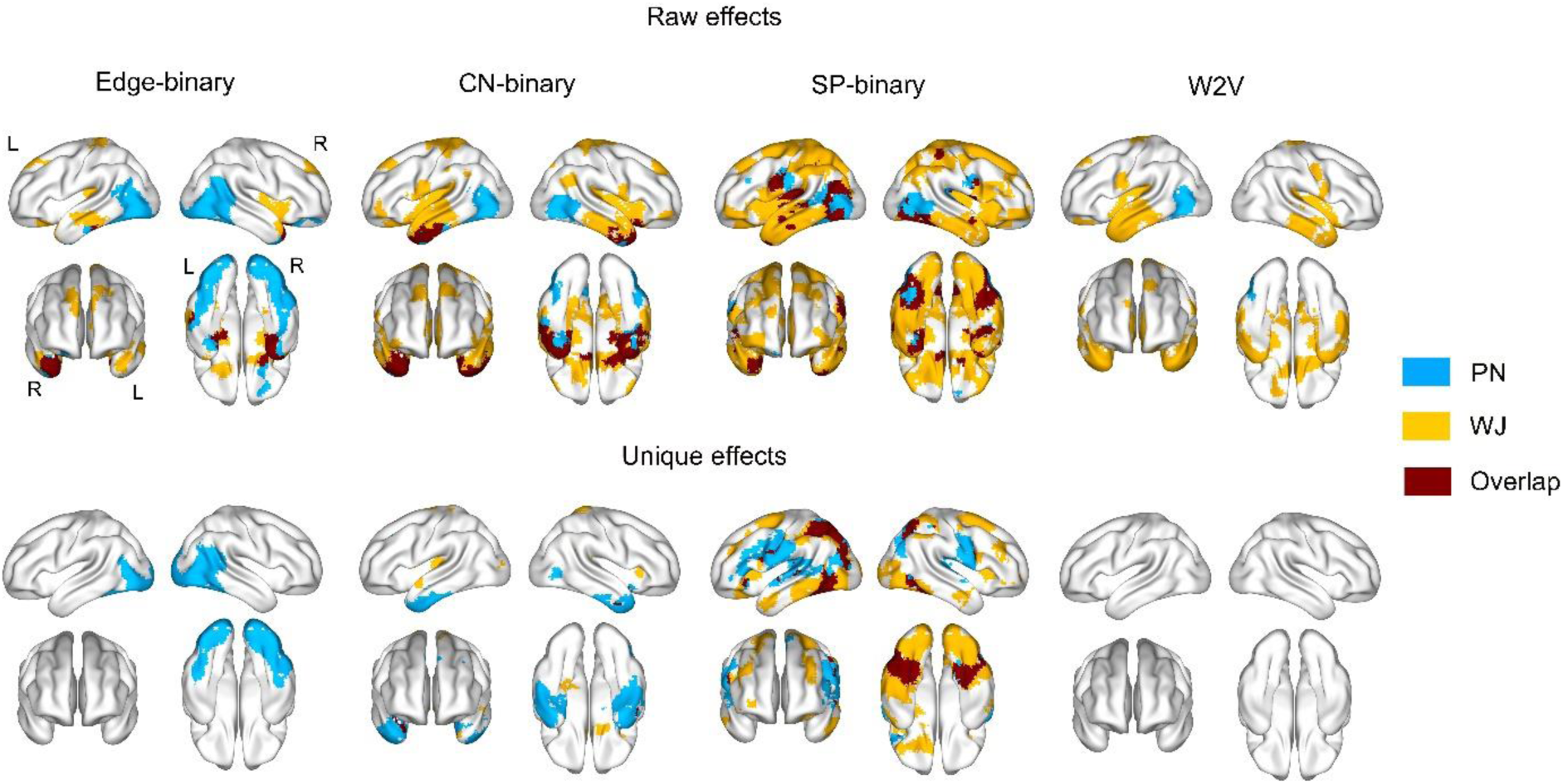
Overlap results of searchlight RSA of language computation models across two experiments using binary graph representation. Following the analysis procedure in Fig. 2a, searchlight mappings were conducted using binary versions of graph-related measures (the edges with PPMI greater than 0 were assigned to 1 and other edges to 0). The results were largely similar, except that the unique effects of binary shortest path revealed additional parietal cortex (uncorrected voxelwise *p* < 0.005, cluster size > 20 in each type of experiment).

**Fig. S6.**
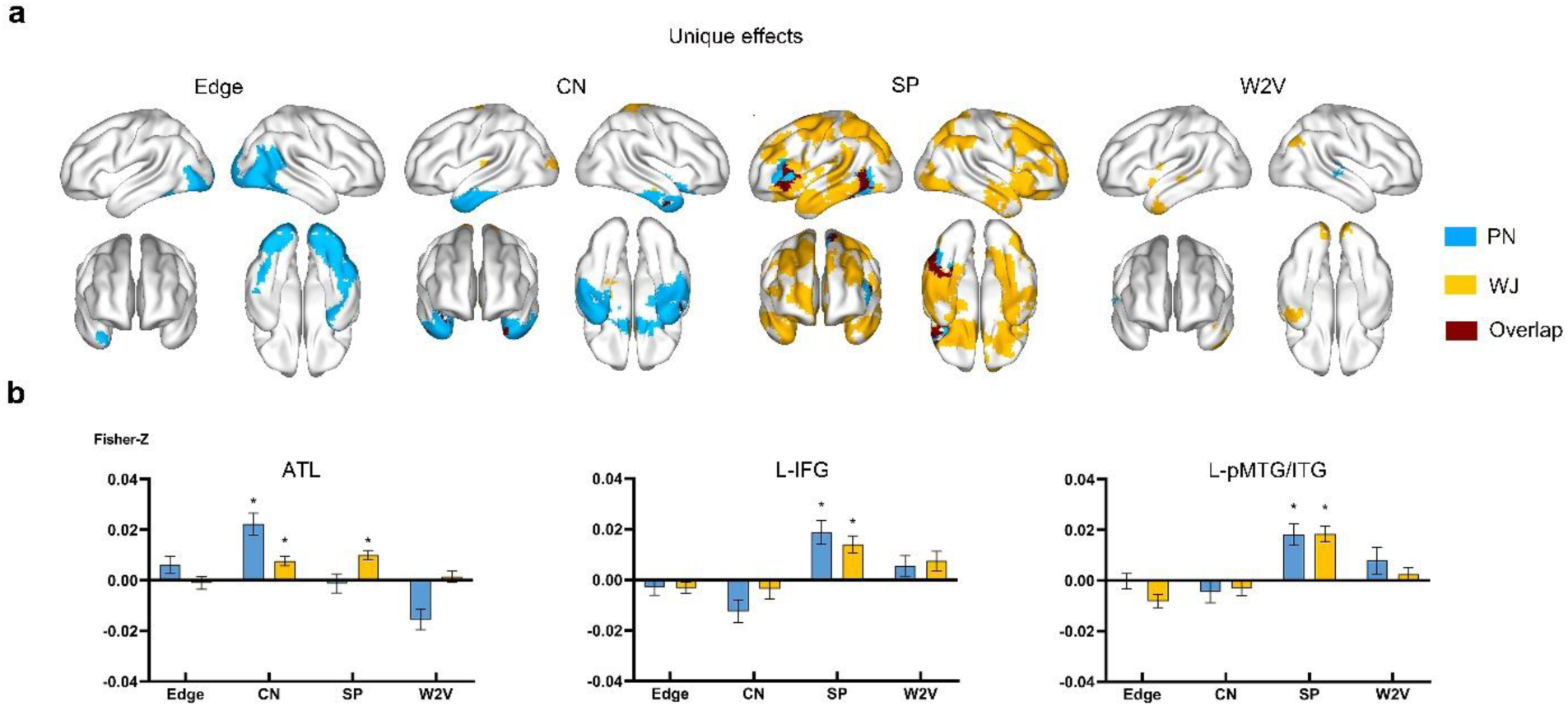
Overlap results of searchlight RSA of language computational models across two experiments using another word2vec vector representation. Following the analysis procedure in Fig. 2a, searchlight mappings were conducted using words’ cosine distance from a word2vec model that was trained on the same corpus on which the edge and graph-related measures were obtained (see Main text). The parameter setting of the word2vec model was similar to the pre-trained word2vec model adopted in the main analysis: skip-gram with negative sampling, dynamic window size = 5, subsampling rate = 10^-5^, negative sample number = 20, context distribution smoothing = 0.75, dimension = 300). (a) Overlap of whole brain searchlight maps from the unique effects of language models across experiments (uncorrected voxelwise *p* < 0.005, cluster size > 20 in each type of experiment). (b) ROI results using ROIs from Fig.2c. The results in Fig. 2c was replicated, indicating it was not the corpus size or content selection that contribute to the observed neural difference between graph and vector embedding models.

**Table S1.** Words (Chinese words and English translations) used in the word familiarity judgment experiment.

| Stimuli |  |
| --- | --- |
| Animals (32): | 袋鼠(kangaroo); 猫(cat); 猴子(monkey); 公鸡(cock); 骆驼(camel); 河马(hippo); 老鼠(mouse); 奶牛(cow); 螃蟹(crab); 大象(elephant); 天鹅(swan); 松鼠(squirrel); 斑马(zebra); 蜘蛛(spider); 狮子(lion); 猫头鹰(owl); 青蛙(frog); 海豚(dolphin); 熊猫(panda); 企鹅(penguin); 鸵鸟(ostrich); 孔雀(peacock); 蚂蚁(ant); 熊(bear); 兔子(rabbit); 老虎(tiger); 鳄鱼(alligator); 蛇(snake); 乌龟(turtle); 蜜蜂(bee); 蜻蜓(dragonfly); 蜗牛(snail) |
| Small Manipulable Artifacts (35) | 锤子(hammer); 筷子(chopstick); 斧头(ax); 剪刀(scissors); 扇子(shanzi); 锯子(saw); 勺子(spoon); 扫帚(broom); 钓竿(fishpole); 算盘(abacus); 扳手(wrench); 刷子(paintbrush); 熨斗(iron); 吸管(straw); 纽扣(button); 刀(knife); 曲别针(paperclip); 钥匙(key); 夹子(clip); 火柴(matchstick); 钢笔(pen); 铲子(shovel); 拐杖(crutch); 螺丝刀(screwdriver); 注射器(syringe); 遥控器(remote control); 相机(camera); 信封(envelope); 手表(watch); 电吹风(hair_dryer); 书包(backpack); 眼镜(glasses); 茶壶(teapot); 伞(umbrella); 项链(necklace) |
| Large Non-manipulable Artifacts (28): | 帐篷(tent); 碑(monument); 桶(barrel); 喇叭(speaker); 黑板(blackboard); 锚(anchor); 书架(bookcase); 火车(train); 亭子(pavilion); 滑梯(slide); 电梯(escalator); 暖气片(heater); 降落伞(parachute); 栅栏(fence); 城堡(castle); 站牌(bus_stop); 烟囱(chimney); 坦克(tank); 浴缸(bathtub); 帆船(yawl); 沙发(couch); 桌子(desk); 椅子(chair); 柜子(cabinet); 床(bed); 冰箱(fridge); 洗衣机(washer); 电脑(computer) |

**Table S2.**
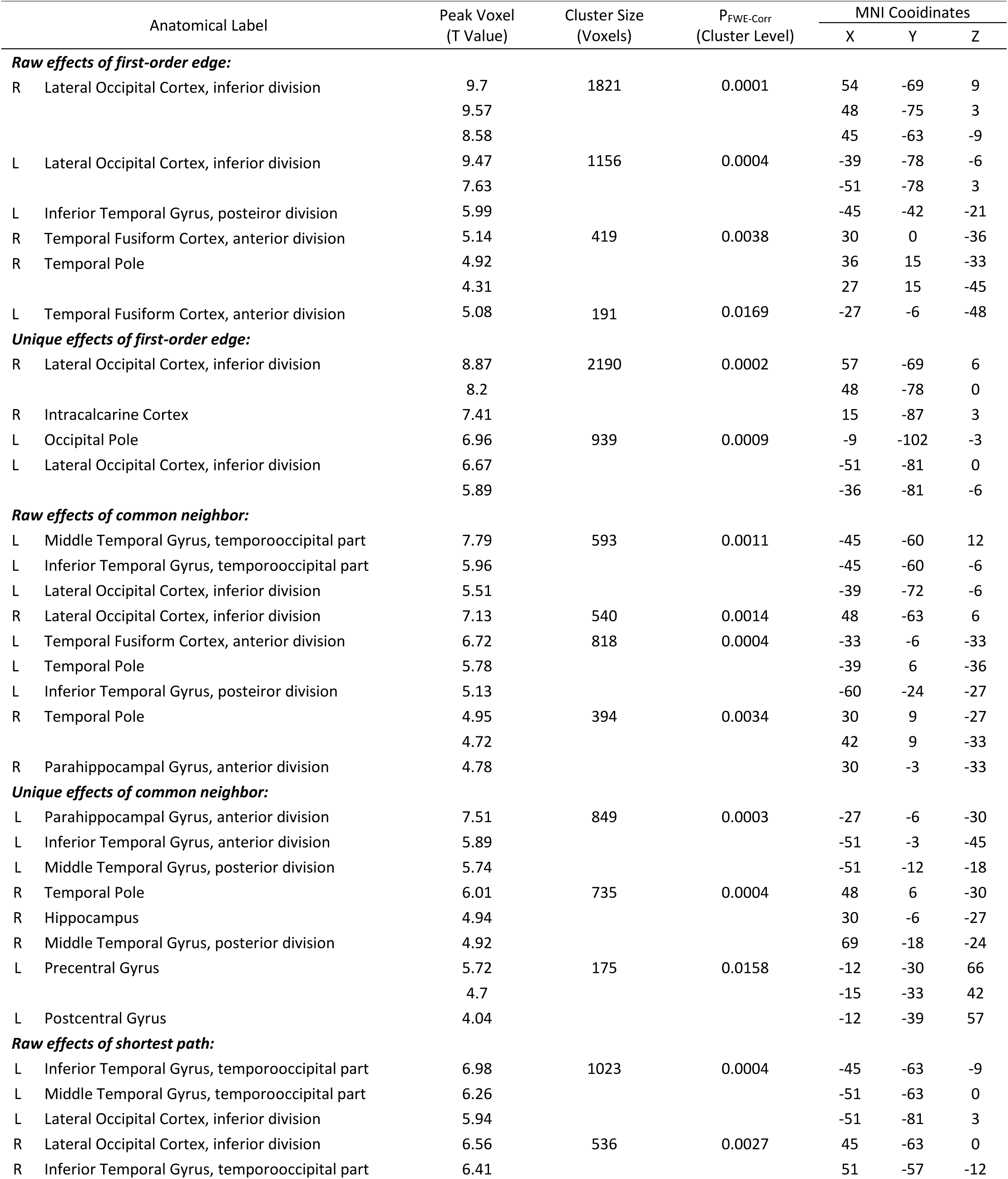
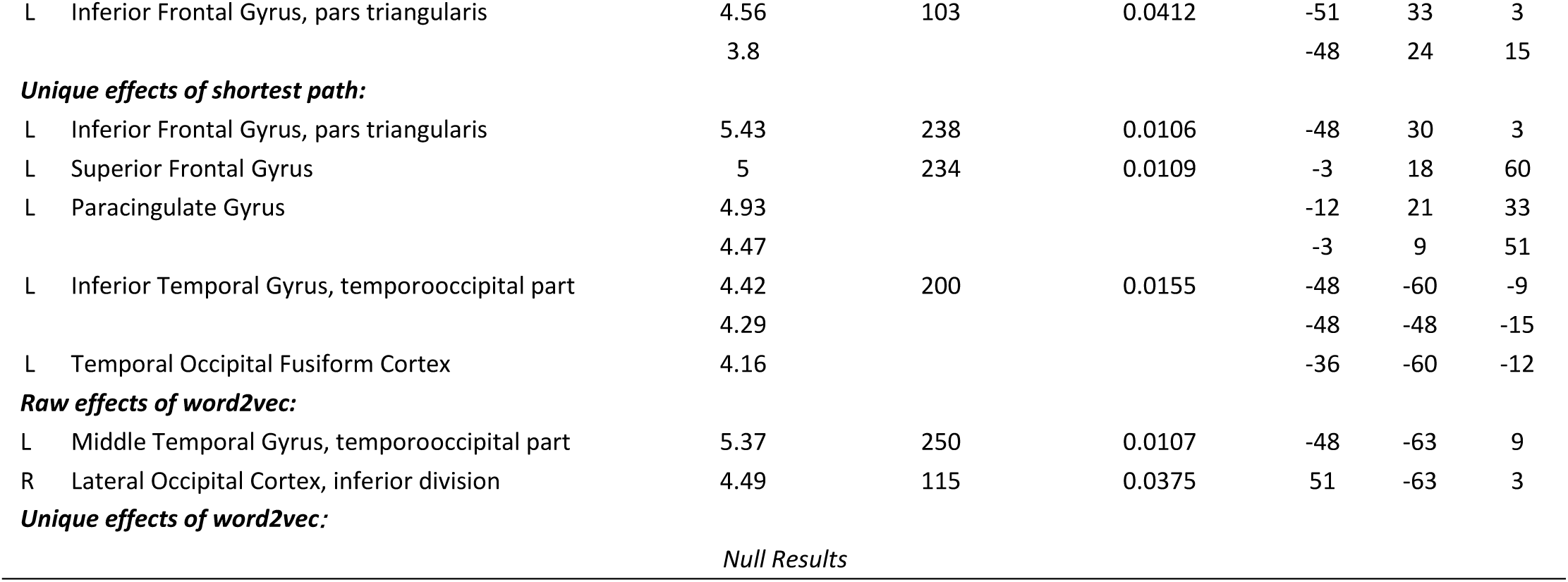
Whole brain searchlight RSA results of language statistical models in the oral picture naming experiment. Results surviving primary voxelwise *p* < .001, one-tailed, cluster-level FWE- corrected *p* < .05 in the whole brain searchlight analysis are presented. Regions are labeled according

**Table S3.**
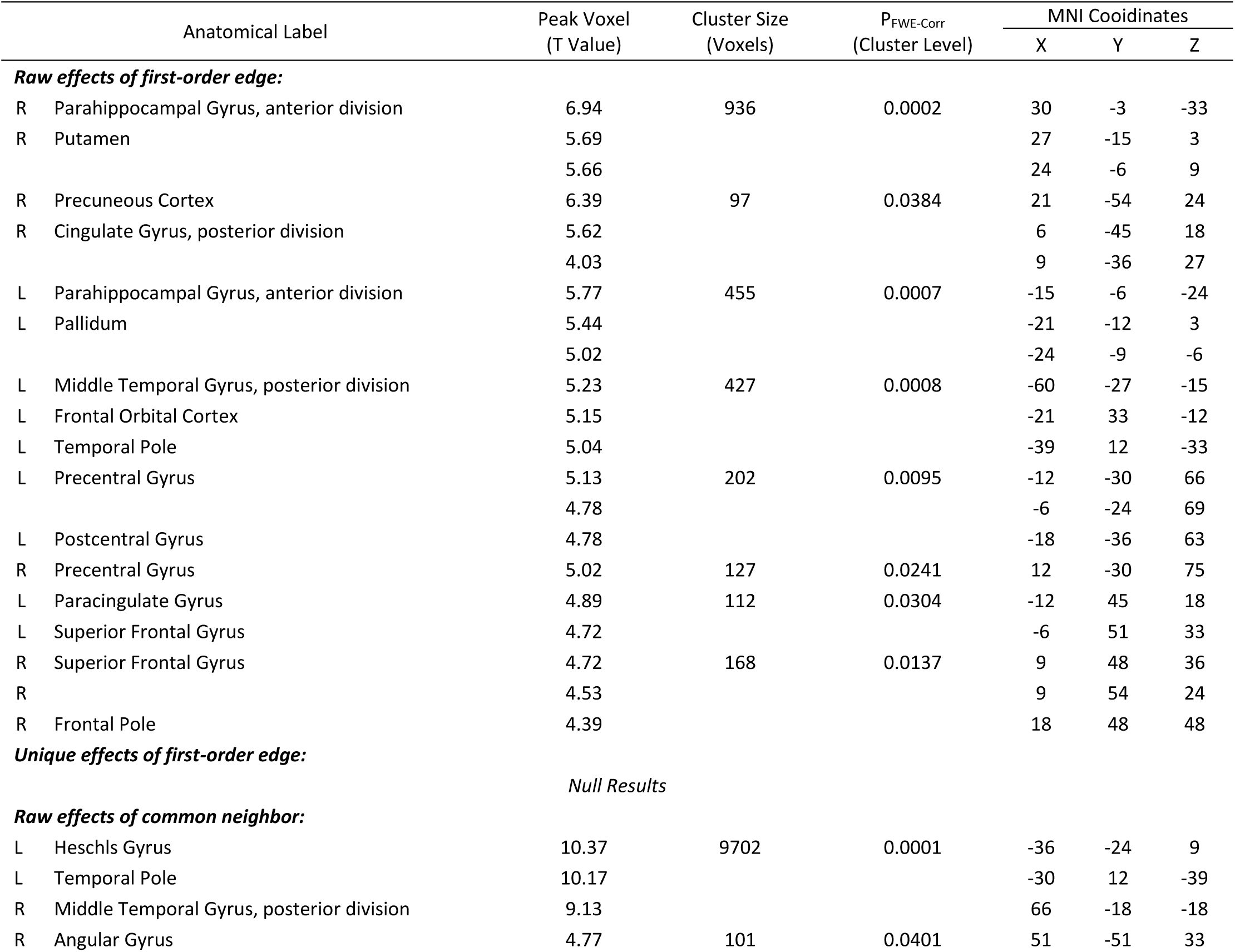
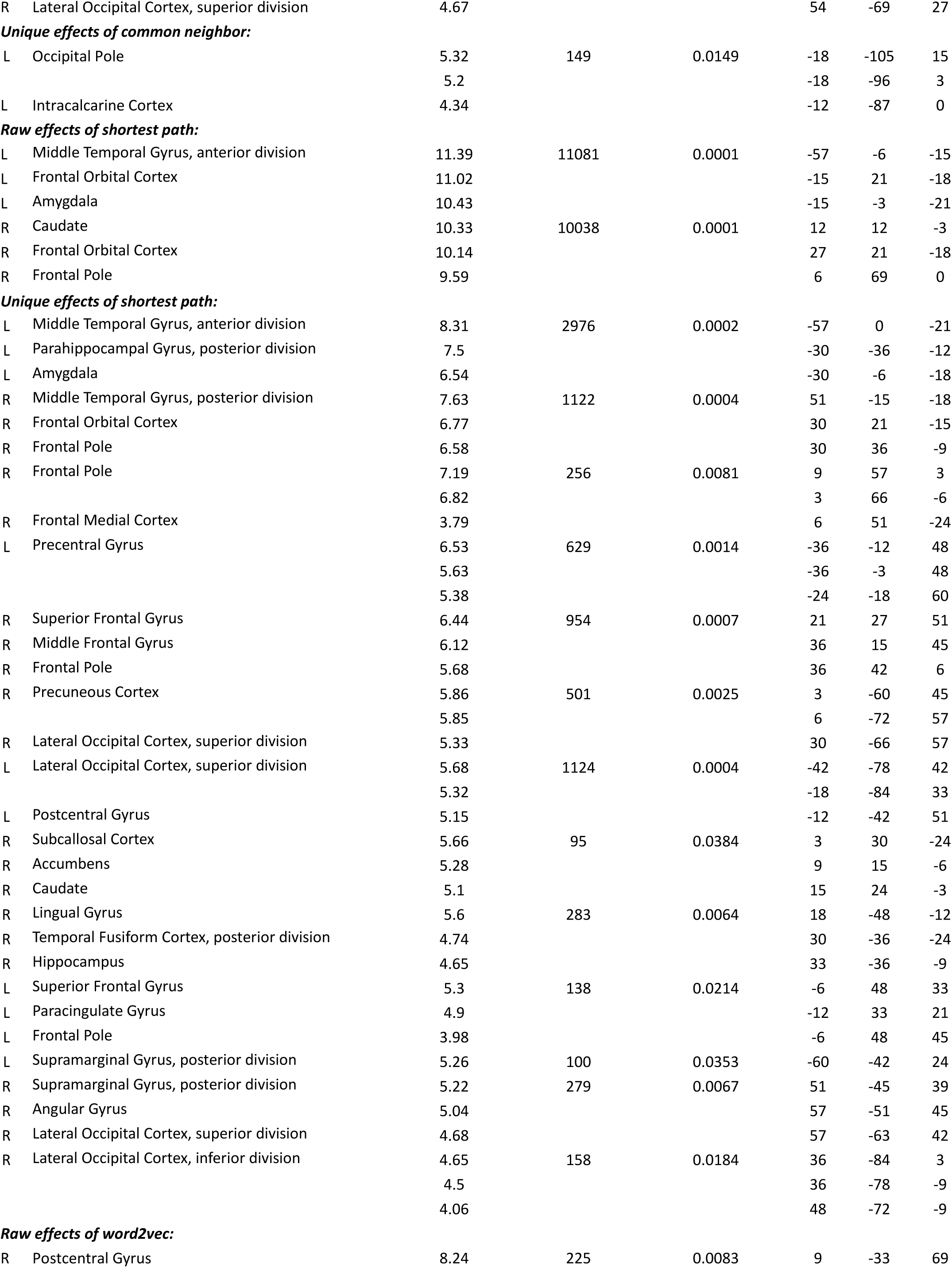
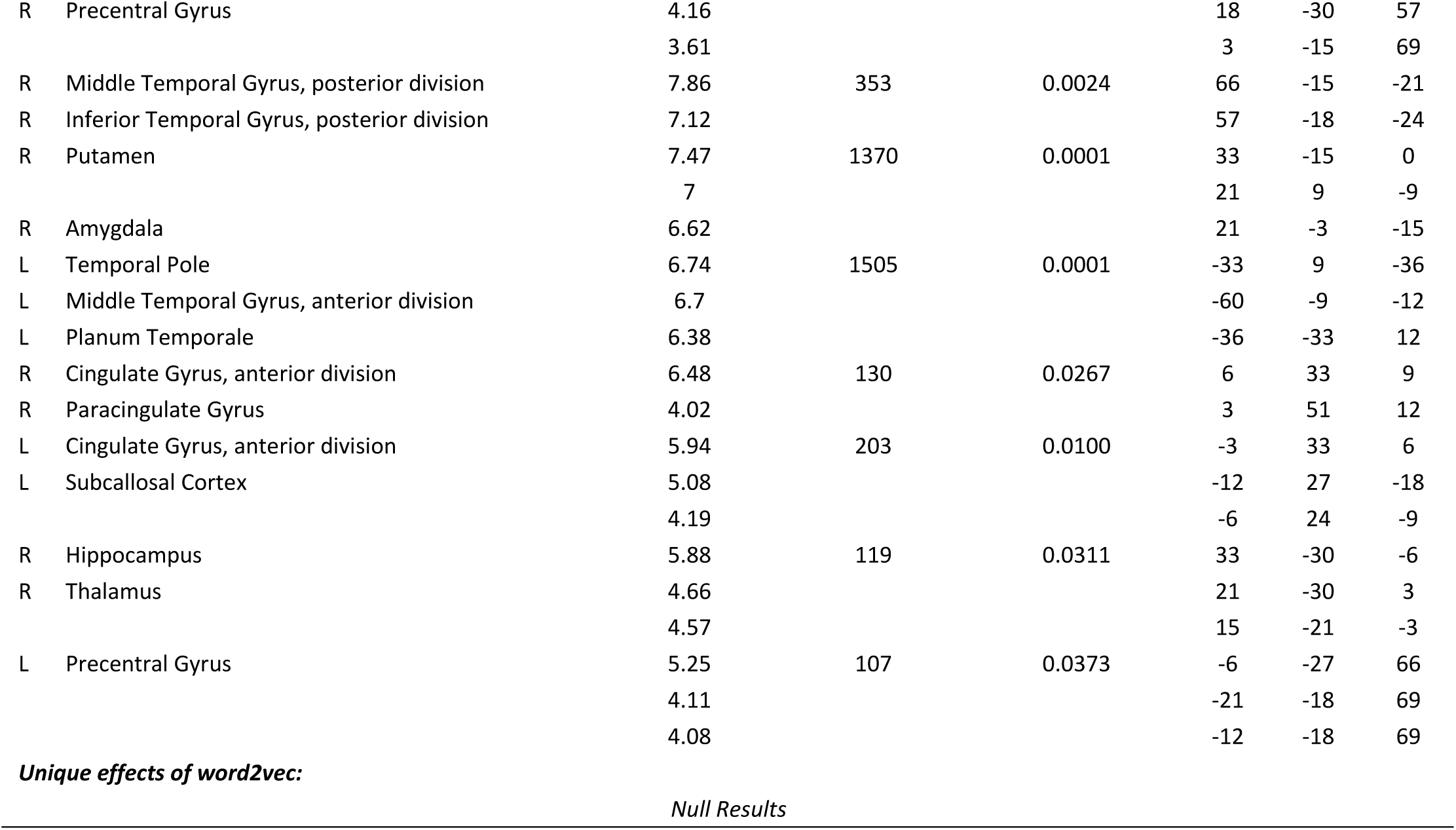
Whole brain searchlight RSA results of language statistical models in the word familiarity judgment experiment. Results surviving primary voxelwise *p* < .001, one-tailed, cluster-level FWE-corrected *p* < .05 in the whole brain searchlight analysis are presented. Regions are labeled according to the Harvard-Oxford cortical and subcortical atlas.

